# Functional redundancy secures resilience of chain elongation communities upon pH shifts in closed bioreactor ecosystems

**DOI:** 10.1101/2022.11.12.516197

**Authors:** Bin Liu, Heike Sträuber, Florian Centler, Hauke Harms, Ulisses Nunes da Rocha, Sabine Kleinsteuber

## Abstract

For anaerobic mixed cultures performing microbial chain elongation, it is unclear how pH alterations affect the abundance of key players, microbial interactions and community functioning in terms of medium-chain carboxylate yields. We explored pH effects on mixed cultures enriched in continuous anaerobic bioreactors representing closed model ecosystems. Increasing the pH from 5.5 to 6.0 caused fluctuations in community composition and yields of *n*-butyrate, *n*-caproate and *n*-caprylate. Further pH increase to 6.5 restored previous yield values while the community entered a new state characterized by lower diversity and evenness but apparently higher richness, indicating the presence of species below the detection threshold in the previous state. We applied Aitchison PCA clustering, linear mixed-effects models and random forest classification on our datasets. Different pH preferences of two key chain elongation species – one *Clostridium* IV species related to *Ruminococcaceae* bacterium CPB6, and one *Clostridium sensu stricto* species related to *Clostridium luticellarii* – were determined. Based on relative abundances measured by 16S rRNA amplicon sequencing, network analysis revealed positive correlations of *Clostridium* IV with lactic acid bacteria, which switched from *Olsenella* to *Lactobacillus* along the pH increase, illustrating the plasticity of the food web in chain elongation communities. The pH increase induced dramatic shifts in community composition whereas process performance (in terms of product range and yields) returned to the previous state. Despite long-term cultivation in closed systems over the pH shift experiment, the communities retained functional redundancy in fermentation pathways, reflected by the emergence of rare species and concomitant recovery of chain elongation functions.

## INTRODUCTION

Microbial ecologists aim to understand the main environmental factors driving the processes of microbial community assembly and functioning [1–3]. Ecological selection exerted by abiotic and biotic factors influences the growth rates of community members and their interactions, thereby determining the composition and functioning of microbial communities [4–7]. In engineered systems, pH is a key parameter shaping microbial communities and steering them towards specific functions [8–12].

Here, we explored the effect of pH shifts on mixed cultures previously enriched under well-controlled abiotic conditions in anaerobic bioreactors as model ecosystems [13, 14]. Even without continued inoculation, such closed systems are relatively complex regarding microbial interactions and metabolic processes. Enrichment cultures can maintain their functional stability by self-assembly, which appears challenging for designing synthetic communities, due to the lack of knowledge required to rationally engineer stable microbial interactions [15]. In our model ecosystems, we focused on the process of microbial chain elongation (CE) to produce the carboxylates *n*-butyrate (C4), *n*-caproate (C6) and *n-* caprylate (C8) from xylan and lactate [13]. These model substrates simulate the feedstock conditions in anaerobic fermentation fed with ensiled plant biomass [16]. Lactate-based CE coupled with *in situ* lactate formation holds promise to valorize organic wastes or biomass residues within the carboxylate platform [17]. Efficient and stable CE processes rely on trophic relationships among community members with diverse metabolic functions forming complementary and parallel pathways in a food web [13].

Next-generation sequencing (e.g., 16S rRNA amplicon sequencing) allows capturing the dynamics of entire communities with high phylogenetic resolution over long-term experiments [7], though there are some methodological limitations, such as PCR biases [18]. Amplicon sequencing data (e.g., amplicon sequencing variants – ASV) only provide proportions. Considering the compositionality of such datasets that contain the relationship information between the parts, approaches that usually start with a log-ratio transformation were developed to avoid the common pitfalls in analyzing compositional data [19–21]. For correlation analysis, association network algorithms are commonly applied, inferring non-random co-occurrence patterns between community members and assessing microbial responses to environmental changes. In this study, standard microbiome analysis and compositional data analysis were implemented to achieve statistically more robust results.

Besides pH, time is an essential component in long-term experimental studies. We categorized time as another factor to emphasize the effect of time (of pH shift) on community dynamics, which reflects the ecological memory in our ecosystems. Suitably clocked sampling with replicates over long experimental times gives insight into the stability of microbial communities and their response to and recovery from perturbations [9, 22, 23]. Linear mixed-effects models (LME) and variations thereof are commonly used for modelling time-resolved 16S rRNA amplicon sequencing data, thereby identifying temporal microbial interaction patterns [24, 25]. We hypothesized that the pH value predominantly determines the assembly of CE reactor microbiomes, but the impact of time needs to be disentangled by applying LME. The temporal patterns of identified taxa are crucial to understand their roles in CE functions being inferred from the correlation with measured process parameters. Feature selection using random forest classification was performed to denote bioindicators of pH changes. Subsequently, the genetic potential of these bioindicators was investigated by functional annotation of metagenome-assembled genomes (MAGs) [14]. As for CE, it is still unclear how the different microorganisms interact and what conditions they thrive in. In this context, pH can be a critical parameter that affects these relationships and ultimately the end products of CE. Our study focused on the effects of pH increase considering three aspects: (i) the identity and abundance of key players of lactate-based CE, (ii) the effect on microbial interactions, and (iii) the functional resilience of the CE reactor microbiome.

## MATERIALS AND METHODS

### Reactor operation and sampling

A microbial community was enriched in a 1-l bioreactor (BIOSTAT^®^ A plus, Sartorius, Göttingen, Germany), inoculated with broth from a former study [16] and fed with mineral medium containing xylan and lactate over 150 days [13]. The enriched community producing C4, C6 and C8 was further selected by reducing the hydraulic retention time (HRT) in two parallel BIOSTAT bioreactors (A and B) for almost one year [14]. The pH was kept at 5.5 in both periods. Here, we tested the effect of pH increase with a fixed HRT of 4 d. Before starting the experiment, the microbial communities of bioreactors A and B were equally distributed by pumping the content from A to B and back while maintaining anoxic conditions.

The reactor configuration was similar as reported before [13], with both bioreactors operated at 38 ± 1°C, constantly stirred at 150 rpm and the pH automatically controlled with 5 M NaOH. For daily feeding, 2.94 g lactate and 2.50 g water-soluble xylan were supplied in 0.25 l mineral medium [13]. The same volume of liquid effluent was harvested daily from the reactors before feeding. The starting pH was 5.5 for both bioreactors. After 42 days, we increased the pH of bioreactor A to 6.0 and further to 6.5 from day 112 to day 238. To consider the effect of time on community assembly, a different temporal scheme of pH increase was applied in reactor B (pH 5.5: day 0-144, pH 6.0: day 145-214, pH 6.5: day 215-238).

Reactor headspace and effluent were sampled twice per week. The effluent was centrifuged and the supernatant was used for measuring concentrations of xylan, carboxylates and alcohols [13]. Optical density (OD) at 600 nm of the effluent was measured before centrifugation. Pelleted cells were stored at −20°C for DNA-based community analysis [13].

### Analytical methods

Daily gas production was monitored as described previously [26]. Gas composition was determined in triplicate for H_2_, CO_2_, N_2_, CH4 and O_2_ by gas chromatography [27]. Concentrations of carboxylates and alcohols were analyzed in triplicate by gas chromatography, and xylan was measured by a modified dinitrosalicylic acid reagent method [13]. At the beginning and the end of the experiment, cell mass concentration was calculated from OD values correlated with cell dry mass [13], with mean correlation coefficients of 1 OD_600_ = 0.641 g l^-1^ for bioreactor A and 1 OD_600_ = 0.632 g l^-1^ for bioreactor B.

Total DNA was isolated from frozen cell pellets using the NucleoSpin Microbial DNA Kit (Macherey-Nagel, Düren, Germany). Methods for DNA quality control and quantification were reported previously [28]. 16S rRNA genes were PCR-amplified using primers 341f and 785r [29] and sequenced on the Illumina Miseq platform (Miseq Reagent Kit v3, 2×300 bp; Illumina, San Diego, CA, USA).

### Microbiome data analysis

The QIIME 2 v2020.2 pipeline [30] with DADA2 plugin [31] was applied to demultiplex sequences, filter phiX reads, denoising, merging read pairs, trimming and removing chimeras. A total of 6 855 572 sequences ranging from 21 389 to 66 272 read pairs per sample were obtained, with a median of 50 439 in 136 samples. A feature table was created indicating the frequency of each ASV clustered at 100% identity. ASVs with frequencies >2 in at least three samples were kept for further analyses. Taxonomy was assigned with a naïve Bayes classifier trained on the database MiDAS 2.1 [32] and curated with the RDP Classifier 2.2 [33] (confidence threshold: 80%). The filtered ASV table was rarefied to 21 389 reads for downstream analyses (rarefaction curves reached the plateau, Figure S1). A total of 97 unique ASVs remained.

α-Diversity based on rarefied ASV data was evaluated by calculating diversity, evenness and richness [34]. The indices of order one (^1^D and ^1^E) quantify the diversity and evenness by weighting all ASVs equally, while the indices of order two (^2^D and ^2^E) give more weight to the dominant ASVs. Considering the compositional nature of amplicon sequencing data [18], we analyzed the data with standard approaches and their compositional replacements. For dissimilarities in community composition (β-diversity), we used Bray-Curtis distance-based principle coordinate analysis (PCoA) [35] and Aitchison principal component analysis (PCA) via DEICODE, which is robust to data sparsity [19]. The QIIME 2 plugin Qurro [36] was used to visualize and explore feature rankings in the produced DEICODE biplot. PERMANOVA (“adonis” function in R vegan package, v2.5.6; 999 permutations) [20] was used for statistical analyses of β-diversity, with *P* values adjusted according to the false discovery rate controlling procedure introduced by Benjamini and Hochberg [37].

### Statistical analysis of effects of pH increase on reactor microbiota time series

A redundancy analysis-based variation partitioning analysis (VPA) was used to quantify the relative contribution of individual process parameters (pH and time) and their interactive effects on temporal variation in community composition. VPA was performed using the “varpart” function in the R package vegan. We performed a partial Mantel test for each process parameter to examine its correlation with community composition represented by Aitchison and Bray-Curtis distances, independent of time (9999 permutations) using vegan.

The QIIME 2 plugin q2-longitudinal with default settings was used to construct the LME for regression analyses involving dependent data [25]. Random intercept models (REML method) were used to track longitudinal changes of metrics including α- and β-diversity and ASV abundances. In brief, pH and time were designated as fixed effects and bioreactor as a random effect, whereas values represent samples of a random collection. The response variables are the following metrics: ^1^D, ^2^D, ^1^E, ^2^E, richness, PC1 of Aitchison or Bray-Curtis and ASV abundance.

The Microbial Temporal Variability Linear Mixed Model (MTV-LMM) was used to identify autoregressive taxa and predict their relative abundances at later time points [24]. The model assumes that the temporal changes in relative abundance of ASVs are a time-homogenous high-order Markov process. To select the core time-dependent taxa, MTV-LMM was applied to each individual pH level, which generated a temporal kinship matrix representing the similarity between every pair of normalized ASV abundances (a given time for a given individual) across time. A concept of time-explainability was introduced to quantify the temporal variance explained by the microbial community at previous time points.

### Random forest (RF) classification

Supervised classification of pH levels on community compositions was performed using QIIME 2 q2-sample-classifier with default settings [38]. Rarefied ASV data were used as features to train and test the classifier. First, a nested cross-validation of the RF model was applied to overview the classification of the pH levels for all samples. For model optimization, a second layer of cross validation (outer loop) was incorporated to split the dataset into training and test sets five times, and therefore each sample ended up in a test set once. During each iteration of the outer loop, the training set is split again five times in an inner loop to optimize parameter settings for estimation of that fold. Five different final models were trained, with each sample receiving a predicted value. The overall accuracy was calculated by comparing the predicted values to the true values.

Next, we performed a feature selection by randomly picking 80% of the samples to train an RF classifier, and the remaining 20% of the samples were used to test the classification accuracy of the classifier. *K*-fold cross-validation (*K* = 5) was performed during automatic feature selection and parameter optimization steps to tune the model. As determined by using recursive feature elimination, the most important features that maximized model accuracy were selected. Model accuracy and predictions were based on the classifier that utilized the reduced feature set.

### Network analysis

Co-occurrence networks based on rarefied ASV data and process parameter data were inferred by using FlashWeave v0.16 implemented in Julia [21]. FlashWeave uses the centered log-ratio approach for the correction of compositional microbial abundances and infers direct associations. Three networks were constructed for the three individual pH levels, which featured a correlation coefficient <-0.5 or >0.5. Another network was constructed from the entire data of all pH levels. All networks were visualized in Cytoscape v3.8.0 [39].

## RESULTS

### Fluctuation and recovery of process performance

The pH increase from 5.5 to 6.0 caused fluctuations in fermentation products and lactate concentrations, which were not observed upon further increase to 6.5 (Figure 1). First we applied the pH increase in bioreactor A, which immediately presented an increased C8 concentration (mmol C/l) up to 29.1, corresponding to a yield (C mole product to substrate ratio) of 5.2, and a relatively stable yield of C6 (16.0 ± 1.5 at pH 6.0). Lactate accumulated to a concentration of 147.5, while C4 concentration dropped to 69.1, with a yield of 12.1 (Figure 1a). The pH increase left the fast consumption of xylan unaffected (Figure S2). Afterwards, accumulated lactate was consumed and C4 concentration returned to the previous level with 273.9 on day 95 at pH 6.0. Notably, further pH increase to 6.5 did not result in such fluctuations (Figure 1a). Later, we replicated the pH increase from 5.5 to 6.5 in bioreactor B to confirm the observed effects of pH increase. With longer operation at pH 5.5 for 144 days, comparable fluctuations in concentrations of lactate, C4 and C8 were observed, but with a delay of 38 days after the pH increase to 6.0. Concentrations of lactate, C4, C6 and C8 were relatively stable when bioreactor B was operated at pH 6.5. The pH increase also resulted in fluctuations of daily gas production and gas composition (Figure S3). A general upward trend of cell mass yield at pH 6.5 suggests a facilitating effect of higher pH on the growth of enriched populations (Figure S4).

**Figure 1.**
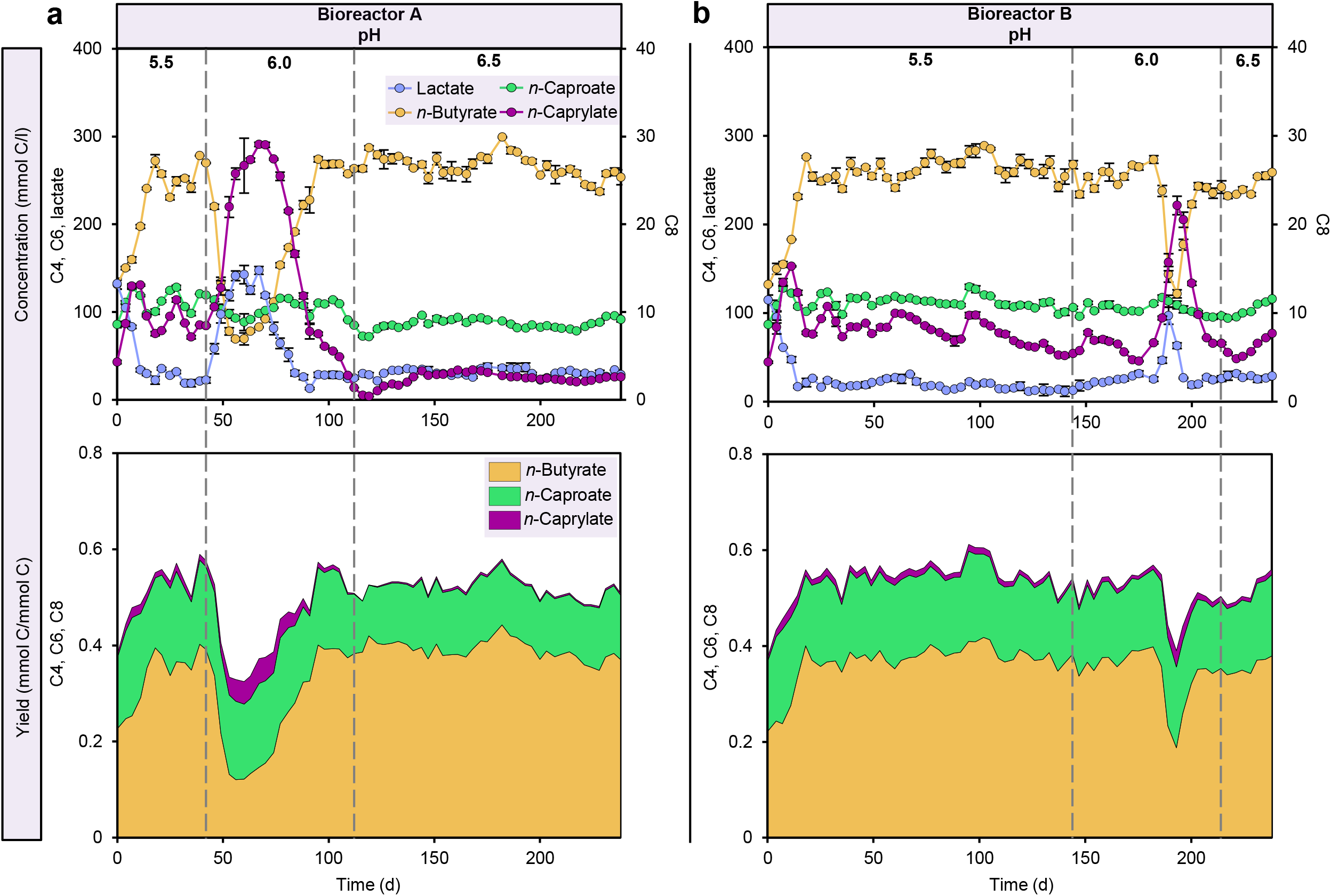
Performance of bioreactors. Concentrations of chain elongation products and lactate, as well as yields of chain elongation products in bioreactors A (**a**) and B (**b**) at three pH levels. Yield is given in C mole product to the transferred substrate ratio. Chain elongation products: C4, *n*-butyrate; C6, *n*-caproate; C8, *n*-caprylate.

### Microbial community shifts and emergence of rare species

After the pH increase, α-diversity metrics showed decreases in diversity (^1^D) and evenness (^1^E), but an increase in richness (Figure 2; similar results for ^2^D and ^2^E shown in Figure S5). We used LME models to test whether these indices were impacted by pH and time (of pH shift), which presents the memory effect on community dynamics. Three separate LME models were fitted to examine ^1^D, ^1^E and richness across pH gradients because the trajectories appeared nearly linear. Diversity was significantly impacted by pH (*P* < 0.001) and time (*P* < 0.001), indicating that diversity was reduced much stronger by pH with a factor of 6.188 than by time with a factor of 0.209 (Table S1). Evenness and richness were also significantly associated with pH and time, although pH exerted much stronger impacts on both indices (Tables S2-S3). As shown in Figure S6, the relative ASV abundances categorized from phylum to genus level varied along the pH gradients, e.g., *Actinomyces* and *Prevotella* became apparent at pH 6.5 along with an increasing abundance of *Clostridium sensu stricto* and decreasing abundances of *Clostridium* IV and *Eubacterium* (Figure S6e).

**Figure 2.**
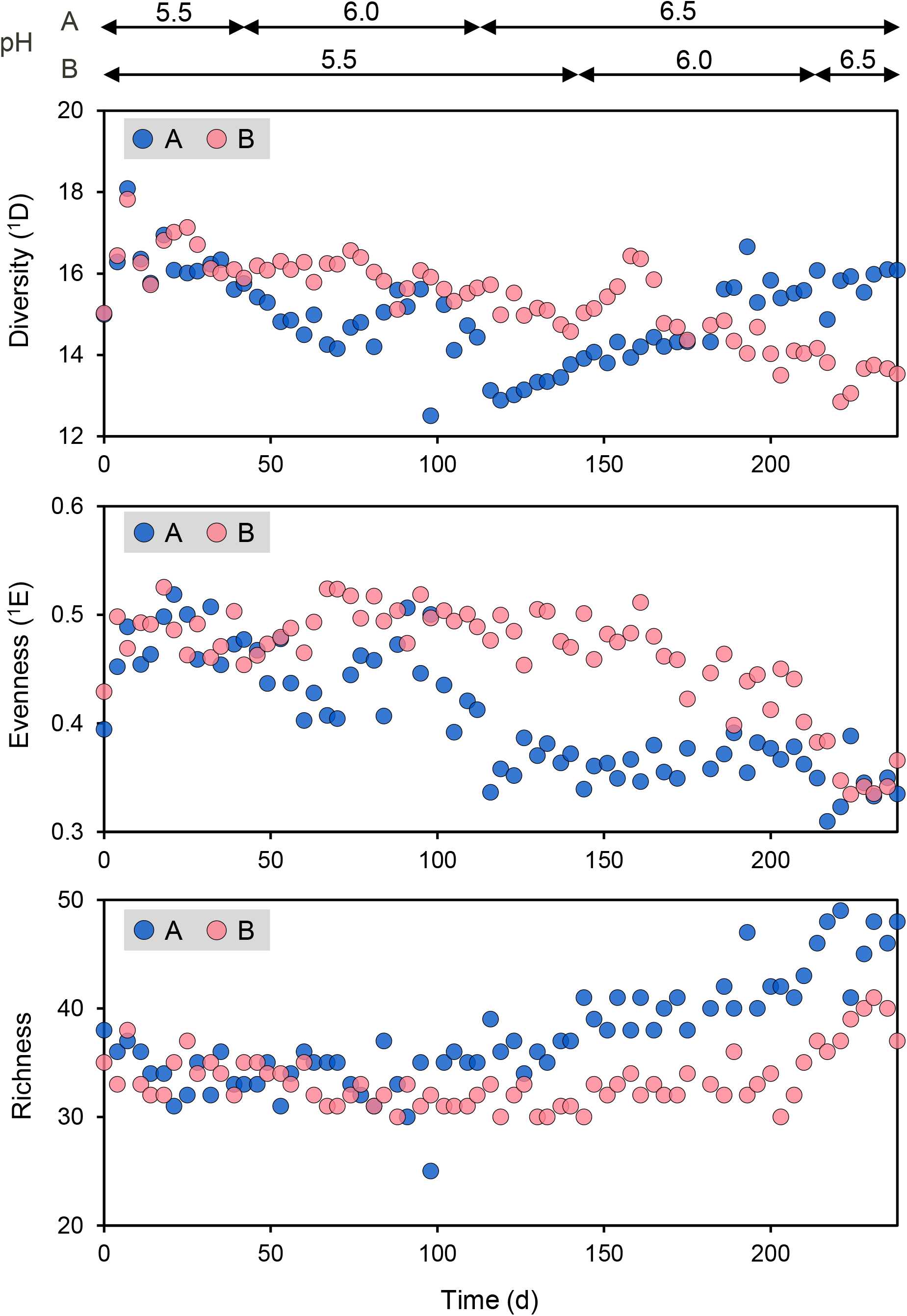
Longitudinal changes in α-diversity at three pH levels. Based on the relative abundances of ASVs, we calculated the α-diversity represented by diversity of order one (^1^D) (**a**), evenness of order one (^1^E) (**b**), and richness (**c**). Diversity and evenness of order one were quantified by weighting all ASVs equally. A and B stand for bioreactors A and B.

β-Diversity analysis revealed that the bacterial communities differed significantly between the three pH levels (PERMANOVA; *P* < 0.001) (Figures 3a, S7). ASVs of *Clostridium* IV, *Oscillibacter, Olsenella* and *Syntrophococcus* were strongly associated with the communities at pH 5.5 and 6.0, while *Clostridium sensu stricto* ASV009 was most strongly associated with the communities at pH 6.5 (Figure 3a). Based on the association with dissimilarities in community composition, *Clostridium* IV ASV008 (lowest ranked taxon) and *Clostridium sensu stricto* ASV009 (highest ranked taxon) correspond to the most influential taxa driving the Aitchison PCA clustering (Figure 3b). After fitting LME models to their dynamics in relative abundance (Figure 3c), results showed that the relative abundance of ASV008 was significantly impacted by pH (*P* < 0.001) and time (*P* = 0.002), whereas only pH (*P* < 0.001) significantly impacted the abundance of ASV009 (Tables S4-S5). In both cases, pH had a much stronger impact than time. By applying LME models, we examined how β-diversity changed over time in each bioreactor (Figure 3d–3e). Results indicated that pH was the most influencing factor, although time had significant effects as well (Tables S6-S7). The partial Mantel test confirmed the impact of pH on the community assembly. We correlated the time-corrected dissimilarities of community composition with pH, and the results show strong, significant correlations based on Aitchison distance (*r_m_* = 0.61, *P* < 0.001) and Bray-Curtis distance (*r_m_* = 0.72, *P* < 0.001) (Table 1). Evaluation of the overall contributions of pH and time by VPA indicated that together they explain 61% of the microbial community variations based on Bray-Curtis (Figure S8). 24% and 3% of the variations were independently explained by pH and time, respectively. These results support those inferred from the LME models.

**Figure 3.**
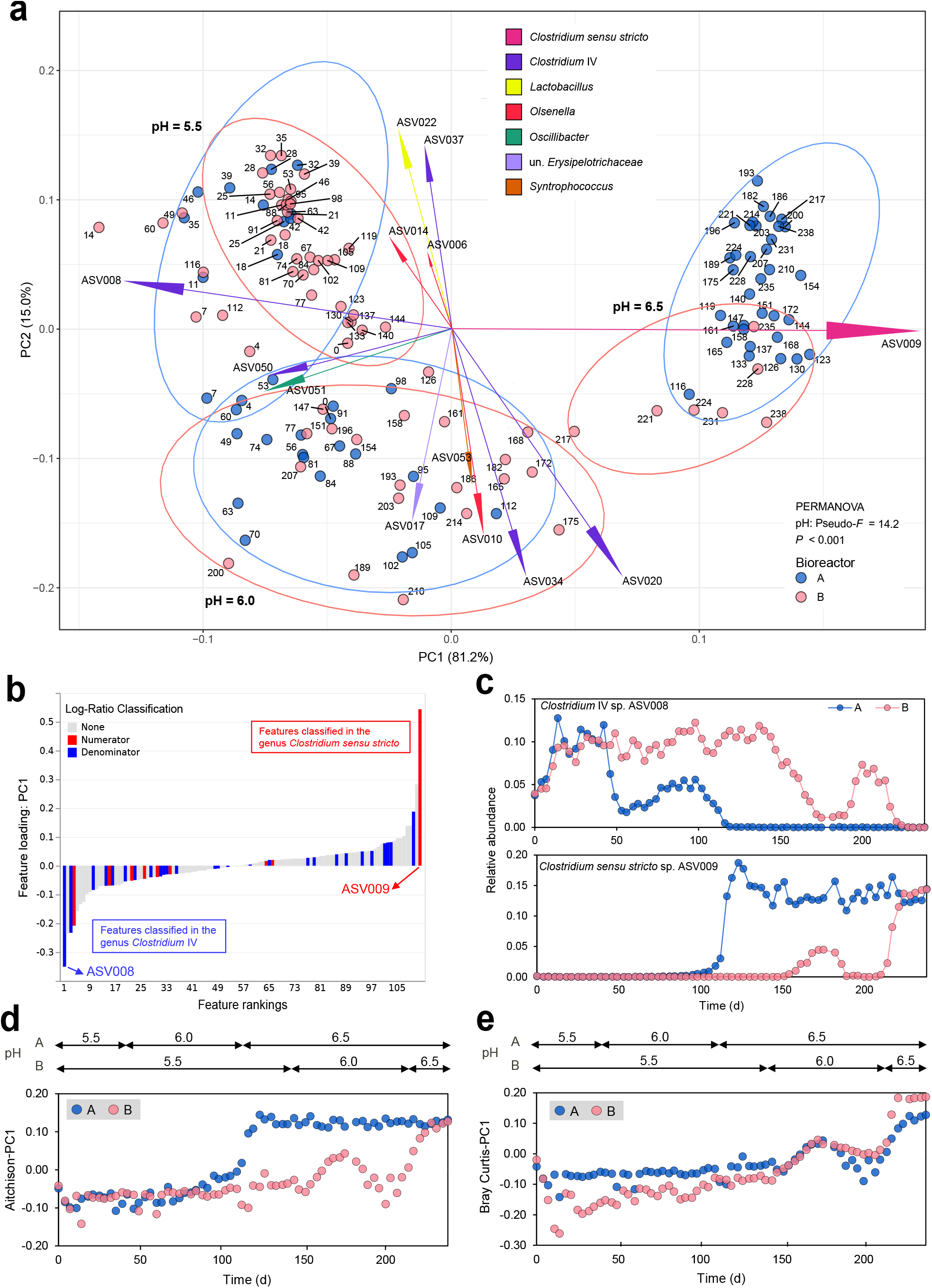
Effects of pH increase and time on bacterial community composition. **a,** Variance-based compositional principal component analysis (PCA) biplot based on Aitchison distance. Dots are named according to sampling days. Ellipses of 95% confidence intervals were added to each individual pH level of the bioreactors. The size of an ASV arrow indicates the strength of the relationship of that ASV to the community composition. ASVs are colored by family. **b,** ASV ranks estimated from Aitchison distance-based PCA (PC1) with *Clostridium* IV and *Clostridium sensu stricto* highlighted. **c,** Longitudinal changes in relative abundances of *Clostridium* IV sp. ASV008 and *Clostridium sensu stricto* sp. ASV009 at three pH levels. **d, e,** Longitudinal changes in β-diversity at three pH levels, based on Aitchison (**d**) and Bray-Curtis (**e**) dissimilarities. A and B stand for bioreactors A and B. un., unclassified.

**Table 1.** Partial Mantel tests showing significant correlations between the time-corrected dissimilarities of microbial community composition and process parameters.

| Process parameter | Aitchison distance |  | Bray-Curtis distance |  |
| --- | --- | --- | --- | --- |
| | $r_m^a$ | $P^b$ | $r_m$ | $P$ |
| pH | 0.61 | < 0.001 | 0.72 | < 0.001 |
| Conc. C2<sup>c</sup> | 0.27 | < 0.001 | 0.18 | < 0.001 |
| Conc. C4 | 0.07 | 0.013 | -0.01 | 0.569 |
| Conc. C6 | 0.29 | < 0.001 | 0.48 | < 0.001 |
| Conc. C8 | 0.25 | < 0.001 | 0.16 | < 0.001 |
| Conc. lactate | 0.02 | 0.258 | 0.01 | 0.401 |
| Conc. biomass | 0.16 | < 0.001 | 0.11 | 0.002 |
| Yield C2 | 0.27 | < 0.001 | 0.15 | < 0.001 |
| Yield C4 | 0.09 | 0.004 | 0.00 | 0.448 |
| Yield C6 | 0.38 | < 0.001 | 0.40 | < 0.001 |
| Yield C8 | 0.22 | < 0.001 | 0.13 | 0.003 |
| Yield biomass | 0.09 | 0.001 | 0.06 | 0.037 |
| O<sub>2</sub> | 0.43 | < 0.001 | 0.44 | < 0.001 |
| CO<sub>2</sub> | 0.14 | < 0.001 | 0.18 | < 0.001 |
| H<sub>2</sub> | 0.19 | < 0.001 | 0.16 | < 0.001 |
| Time | 0.14 | < 0.001 | 0.33 | < 0.001 |

<sup>a</sup> $r_m$ , the correlation coefficient based on partial Mantel test, in which time was controlled. The permutation test compares the original $r_m$ to $r_m$ computed in 9999 random permutations.
<sup>b</sup>The reported $P$ value is one-tailed.
<sup>c</sup>Conc., concentration

### pH bioindicators and time-dependent taxa

Overall, the nested cross-validation of RF classification represented an accuracy of 97.8% in predicting the pH levels for all 136 samples (Figure S9), using ASV data to follow community composition dynamics. We carried out recursive feature elimination with cross-validation; the 18 most important features were selected that gave perfect discrimination between the three pH levels (Figure 4). These ASVs were defined as pH bioindicators, belonging to the genera *Clostridium* IV, *Syntrophococcus, Lactobacillus, Olsenella, Bulleidia, Clostridium sensu stricto, Eubacterium, Lachnospiraceae incertae sedis, Sporanaerobacter* and *Actinomyces* (Figure 4b). Among these pH bioindicators, four increased in abundance while 14 became less abundant along the pH increase. Notably, the most influential ASVs driving the Aitchison PCA clustering were also pH bioindicators, including the abundant taxa *Clostridium* IV ASV008 and *Clostridium sensu stricto* ASV009 (Figure 4b).

**Figure 4.**
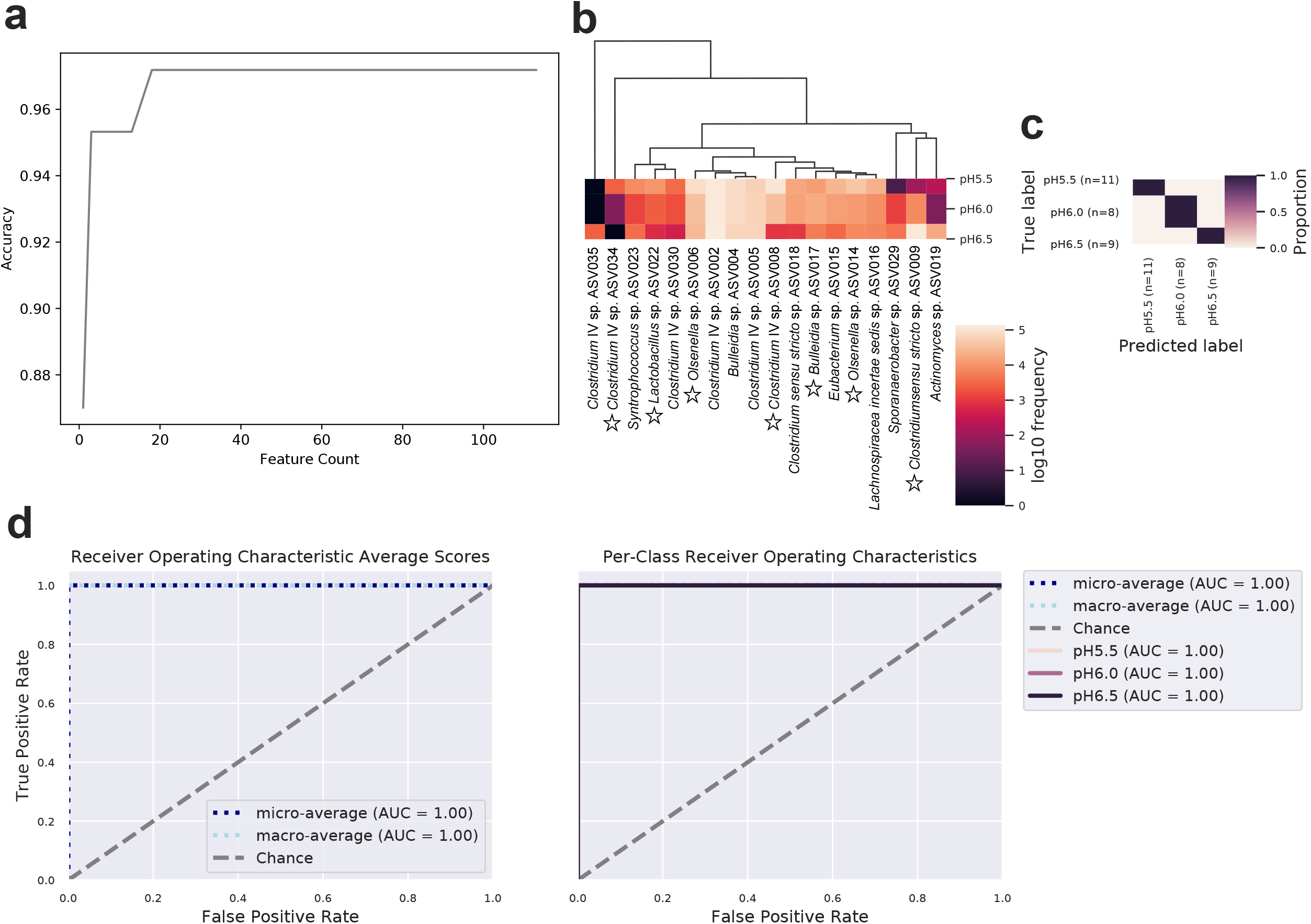
pH bioindicators determined by random forest classification accurately predict the different pH levels. **a,** Recursive feature elimination plot illustrating the model accuracy changes as a function of ASV count. The top-ranked 18 ASVs (pH bioindicators) that maximize accuracy are automatically selected for optimizing the model, based on their mean decrease in Gini scores, according to their ASV abundance distribution, with pH as the response variable. **b,** Heatmap showing dynamics of the mean abundance of pH bioindicators at the different pH levels. ASVs shown in Aitchison PCA biplot are indicated by a star. **c,** Confusion matrix for the optimal classifier of samples at different pH levels. The classifier was trained on the randomly picked 80% of the samples, which was then tested on the remaining 20%. Overall accuracy was calculated by comparing the predicted values with the true values. **d,** Receiver Operating Characteristic (ROC) and Area Under the Curve (AUC) curves representing the accuracy of random forest classification. The ROC curve plots the relationship between the true positive rate and the false positive rate at various threshold settings. The AUC indicates the probability that the classifier ranks a randomly chosen sample of the given class higher than other classes. The random chance is represented as a diagonal line extending from the lower-left to the upper-right corner. To show the ROC curves for each class, average ROC and AUC were calculated. “Micro-averaging” calculates metrics globally by averaging across each sample; hence, class imbalance impacts this metric. “Macro-averaging” gives equal weight to the classification of each sample.

By using MTV-LMM, we identified time-dependent taxa, whose abundance can be predicted based on the previous community composition. In this longitudinal study, 32, 25 and 40 ASVs were predicted to be significantly (*P* < 0.05) affected by the past composition of the community at pH 5.5, 6.0 and 6.5, respectively, with the time-explainability ranging from 17% to 80%, 17% to 83% and 13% to 96%, respectively (Figure S10).

### Microbial interaction patterns

Partial Mantel test showed significant correlations of the community composition with process performance and the changing conditions (Table 1). Consequently, we constructed an overall network and three separate networks for each pH level to discern the succession of microbial interactions and reveal potential metabolic functions. After the pH increase to 6.5, more nodes and edges and higher average clustering coefficient and heterogeneity were found, suggesting that the overall interaction intensity was higher at pH 6.5 (Table S8). In agreement with Aitchison PCA analysis, pH was significantly correlated with pH bioindicators ASV008 and ASV009 (Figure S11). Changes of interaction patterns over pH are shown in Figure 5. At the family level, *Ruminococcaceae* co-occurred with *Lachnospiraceae* and *Erysipelotrichaceae* at all pH levels, while it co-occurred with *Coriobacteriaceae* only at pH 5.5 (e.g., *Clostridium* IV ASV090 with *Olsenella* ASV049). *Ruminococcaceae* also co-occurred with *Lactobacillaceae* at pH 6.0 and 6.5, and with *Actinomycetaceae* only at pH 6.5 (*Clostridium* IV ASV037 with *Actinomyces* ASV019). *Clostridiaceae* 1 co-occurred with *Clostridiales incertae sedis* XI (*Clostridium sensu stricto* ASV009 with *Sporanaerobacter* ASV029) and *Erysipelotrichaceae* only after the pH increase to 6.0. *Erysipelotrichaceae* showed positive correlations with *Lactobacillaceae* at pH 6.0 and 6.5, where its negative correlation with *Coriobacteriaceae* vanished. Notably, the positive correlation of *Erysipelotrichaceae* (*Bulleidia* ASV004) with *Lachnospiraceae* (*Syntrophococcus* ASV001) was not seen at pH 6.0. The positive correlation between C6 yield and *Eubacterium* ASV015 was presented in the overall network and the individual networks of pH 5.5 and pH 6.0, but not in that of pH 6.5 (Figures S11 and 5). In general, stronger correlations (|r| > 0.5) were observed at pH 6.5, including the negative correlation of *Prevotella* ASV041 with *Bulleidia* ASV017.

**Figure 5.**
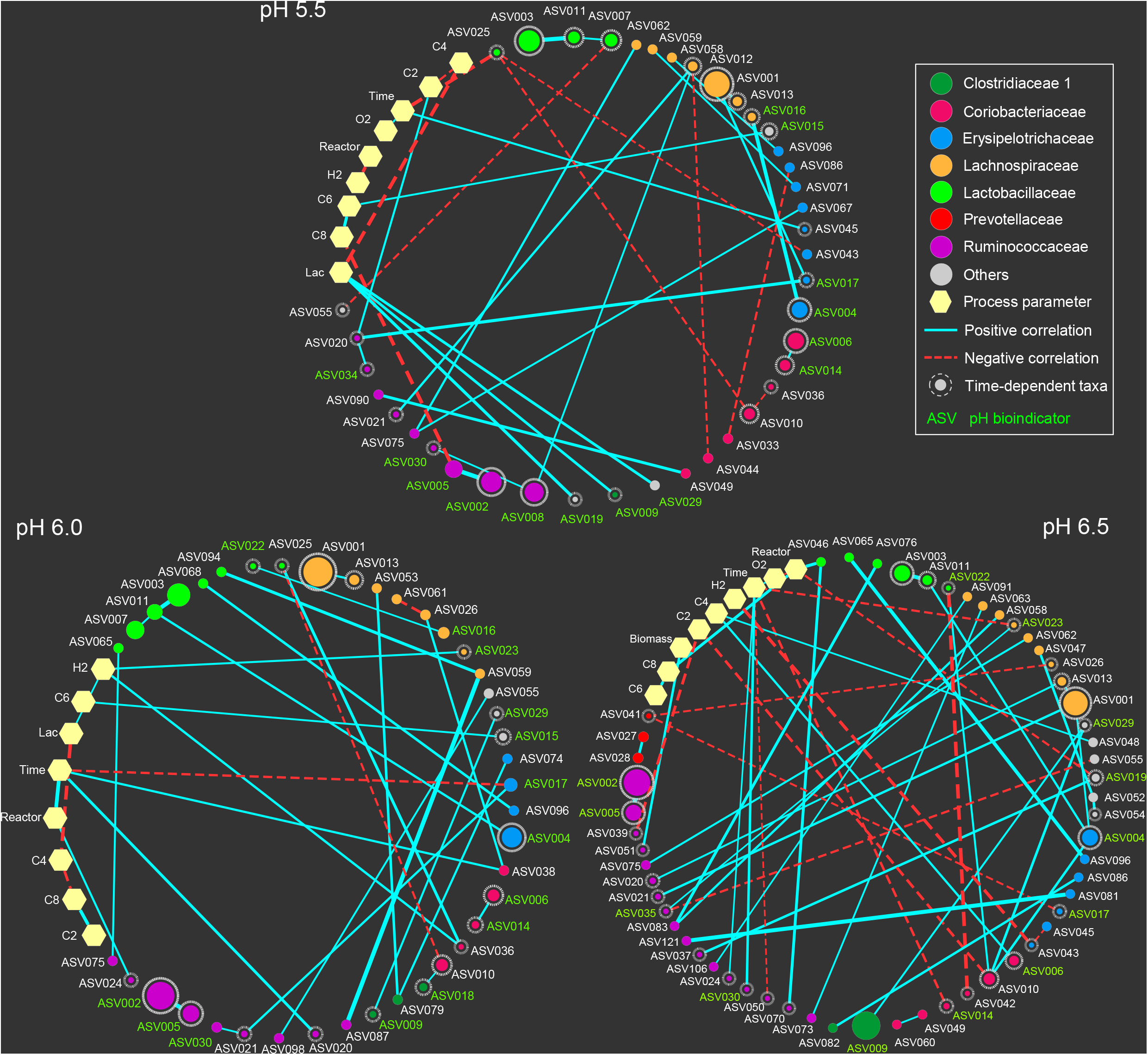
Co-occurrence networks for the three individual pH levels. Edges indicate a coefficient > 0.5 for positive correlations and < −0.5 for negative correlations. Edge thickness reflects the strength of the correlation. The size of each ASV node is proportional to the mean relative abundance over the corresponding pH level. ASV nodes are colored and grouped by family. ASV nodes with grey dashed borders are those time-dependent taxa of each individual pH level, whose abundance can be predicted based on the previous microbial community composition. pH bioindicators identified by random forest classification are shown with green letters. “Others” include the ASVs belonging to families *Eubacteriaceae* (ASV015), *Actinomycetaceae* (ASV019), *Clostridiales incertae sedis* XI (ASV029), *Microbacteriaceae* (ASV048), *Veillonellaceae* (ASV052, ASV054) and *Nocardiaceae* (ASV055). Lac, lactate concentration; C2, acetate yield; C4, *n*-butyrate yield; C6, *n*-caproate yield; C8, *n*-caprylate yield.

## DISCUSSION

### Different pH niches of chain elongation key players *Clostridium* IV and *Clostridium sensu stricto*

MAGs of *Clostridium* IV and *Clostridium sensu stricto* harboring the genetic potential for CE were previously recovered from the enriched communities that served as inoculum for the present study [14; Table S9]. Based on the statistically robust results of Aitchison PCA clustering coupled with LME models and RF classification, a clear conclusion can be drawn: mildly acidic pH values (lower than 6.0) are favorable for *Clostridium* IV while the more neutral pH 6.5 is suitable for *Clostridium sensu stricto*. The corresponding MAGs of *Clostridium* IV ASV008 have 78% average nucleotide identity (ANI) to the lactate-based chain elongator *Ruminococcaceae* bacterium CPB6, which belongs to the family *Acutalibacteraceae* UBA4871 according to the Genome Taxonomy Database [40]). Strain CPB6 was described to prefer mildly acidic pH (5.5 - 6.0) and to suffer from low growth rates and long lag phases at pH values above 6.0 [41]. MAGs corresponding to *Clostridium sensu stricto* ASV009 showed 81% ANI to *Clostridium luticellarii*, which has a pH optimum of 6.5 [42] and CE capability [43–46]. Functional annotation revealed that all genes necessary for lactate oxidation and reverse β-oxidation are present, i.e. these MAGs represent key players of lactate-based CE in our reactor microbiomes [14]. The corresponding ASV008 and ASV009 were identified as time-dependent taxa that are key to understand the community assembly and can be used to characterize the temporal trajectories of the communities. The pH preferences of *Clostridium* IV ASV008 and *Clostridium sensu stricto* ASV009 tied together with concepts in niche theory suggest that different CE bacteria thrive within a defined range of pH values, and outside this range, they are outcompeted by other, better adapted CE species [47]. Due to the distinct growth optima of different populations, alteration of pH is an important tool to shape and control CE reactor microbiomes.

### The pH value as a key determinant of microbial community assembly

Regular and temporally dense sampling with replicates is crucial to capture compositional patterns of communities inferred from time-series data [7, 22]. Microbial interaction is one of the main factors affecting such time-dependent patterns. Given that pH had a much stronger association with community assembly than time, we conclude that pH was the main driver modulating microbial interactions. Our former studies indicated that lactate-based CE driven by *Olsenella* is an essential feature when maintaining the pH at 5.5 [13, 14]. Along with increasing pH, lactic acid bacteria of the genus *Olsenella* cooperating with the chain elongator *Clostridium* IV were replaced by lactic acid bacteria of the genus *Lactobacillus*. Both genera are xylose-fermenting lactate producers according to the functional annotation of their MAGs (Table S9). An enriched community dominated by CE species and *Lactobacillus* was reported in a recent study [48]. Lambrecht et al. [16] suggested inherent benefits of *in situ* lactate formation in CE. The shift in the mutualistic relationship between lactate producers and lactate-consuming chain elongators along the pH gradient revealed the plasticity of the CE microbiota food web. The co-occurrence of phylogenetically closely related taxa may indicate their overlapping metabolic niches, such as the appearance of *Lactobacillus* ASV003 and ASV011, *Syntrophococcus* ASV001 and ASV013, *Clostridium* IV ASV002 and ASV005 at all pH levels.

According to the storage effect, rare species can germinate and become dominant under proper conditions [49, 50]. In this study, the increase in richness can be explained by an abundance shift of some taxa from undetectable to abundant, reflecting the strong inhibition effects of lower pH on these taxa. Although we operated the reactors in a semi-continuous mode, which theoretically leads to the washout of organisms that do not grow fast enough, biofilms might have unintentionally formed, providing a niche for maintaining such rare populations. With the increased number of microbial interactions and increasing interaction intensity strongly coupled to these taxa at higher pH, the factor pH shaping the community assembly was revealed by considering the growth and interactions of community members in such long-term closed systems.

Besides, other effects of pH shifts cannot be ignored. At higher pH, the concentrations of protonated carboxylic acids are lower, which are known growth inhibitors of bacteria including CE community members [11, 51–54]. The longer the chain length, the more toxic the acids are due to the increasing hydrophobicity. On the other hand, the energy gain for CE bacteria is higher with more CE cycles, i.e. longer-chain products. Notably, both bioreactors showed a transient increase of C8 production after rising the pH from 5.5. to 6.0. This might be due to the fact that C8 becomes less toxic at higher pH since a greater share is dissociated, facilitating more CE cycles that lead to C8 formation. Thereafter, C8 production dropped to the previous level, which might be explained by the community shifts caused by the pH increase. There are different terminal enzymes catalysing the reverse β-oxidation and different enzyme complexes involved in energy conservation in CE bacteria, which might have energetic implications for the resulting CE products. Moreover, proton concentration changes and CO_2_-HCO_3_^-^ equilibrium in lactate conversion (3 lactate^-^ + 2 H_2_O → caproate^-^ + 3 HCO_3_^-^ + H^+^ + 2 H_2_) can influence energy release during CE with lactate [55].

### Community changes do not necessarily affect community functioning

We assumed that an increase in pH would induce shifts in the community assembly and consequently community functioning. However, unlike in a complex, open CE system [16], increasing pH had no substantial effects on CE community functioning, i.e., changes in community composition did not necessarily lead to improved carboxylate production during long-term reactor operation. This agrees with the rare associations between ASVs and process parameters in the networks. Without introducing new microorganisms by inoculation, the emergence of rare species indicated high functional redundancy despite the reactors had been operated as long-term closed systems. The reactor performance returned to the previous state after the fluctuation in carboxylate production along pH gradients, suggesting that coexisting rare species can increase functional resilience to environmental disturbances. As mentioned above, the pH shift caused a dramatic but transient increase of C8 yield. How to exploit such disturbance effects for process control needs to be investigated systematically. Keeping functional redundancy in mixed culture processes might be important for biotechnological applications, because parallel pathways of substrate conversion are essential to guarantee the functional stability during perturbation [9, 11, 56]. With regard to the practical implications for mixed-culture processes, our results delineate fundamental differences between long-term enriched microbiomes selected for a specific function and engineered consortia assembled from single species covering all metabolic traits needed for that function. The latter might perform better under stable conditions, whereas naturally selected consortia keeping rare species are more robust under fluctuating conditions and resilient towards perturbations due to their functional redundancy.

## DATA AVAILABILITY

All data described in this study are present in the paper and/or the Supplementary Information. Amplicon sequencing data (ERR4450775 to ERR4450910) have been deposited to the ENA database under study no. PRJEB39808. The MAG sequences used in this study are publicly available in ENA under the accession nos. GCA_903789645, GCA_903789675, GCA_903789585, GCA_903789565, GCA_903789665, GCA_903789475, GCA_903789455, GCA_903789485, GCA_903789575, GCA_903789695 and GCA_903789705.

## SUPPLEMENTARY INFORMATION

Supplementary Figures and Tables: Figures S1-S11 and Tables S1-S9.

## ACKNOWLEDGEMENTS

The authors thank Ute Lohse for skilled technical assistance in molecular analyses and the colleagues from DBFZ Deutsches Biomasseforschungszentrum gGmbH for their technical support in analyses of abiotic parameters. We thank Wanwan Liang from the Centre of Geospatial Analytics at North Carolina State University and Liat Shenhav from Department of Computer Science at University of California Los Angeles for their help with statistics. B.L. was funded by the China Scholarship Council (# 201606350010). F.C., H.S. and S.K. were supported by the BMBF – German Federal Ministry of Education and Research (# 031B0389B, # 01DQ17016 and # 031A317) and the Helmholtz Association (Program Renewable Energies). U.R. was financed by the Helmholtz Young Investigator grant VH-NG-1248 Micro ‘Big Data’.

## AUTHOR CONTRIBUTIONS

B.L., H.S. and S.K. designed the study and the experiments. B.L. performed the experiments, analyzed the process data and sequencing data, and drafted the manuscript. B.L. performed the statistical modeling and machine learning with the help of U.R.. F.C. carried out the network analysis. H.H. contributed to the discussion of the results. All authors contributed to data analysis or interpretation and to the preparation of the manuscript. All authors read and approved the final manuscript.

## COMPETING INTERESTS

The authors declare no competing interests.

## Notes

https://www.ebi.ac.uk/ena/browser/view/PRJEB39808

### Competing Interest Statement

The authors have declared no competing interest.

