## Supplementary Information for "Functional redundancy secures resilience of chain elongation communities upon pH shifts in closed bioreactor ecosystems"

Figure S10. Core time-dependent taxa of individual pH levels……………………..………14

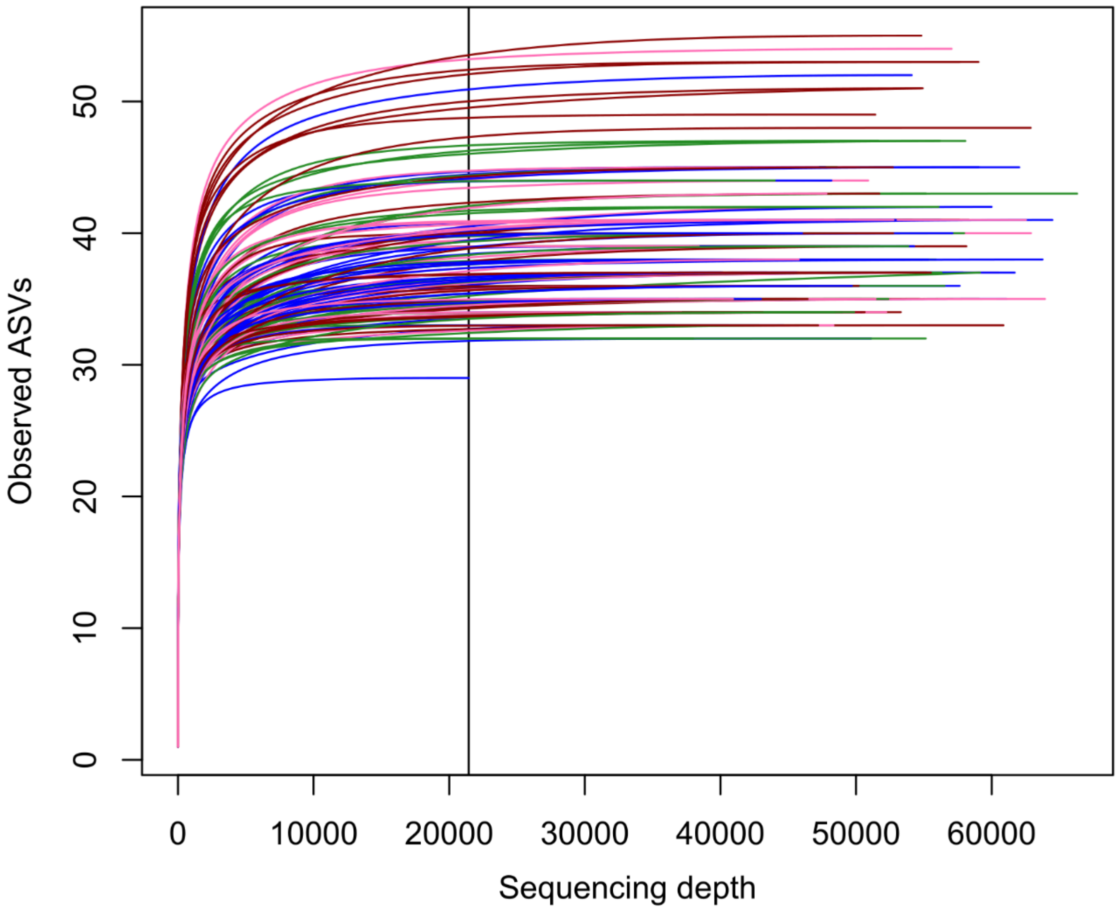

**Figure S1. α-Diversity rarefaction curves.** ASVs of all samples were rarefied to an equal sequencing depth of 21 389 reads. Colors represent the different samples.

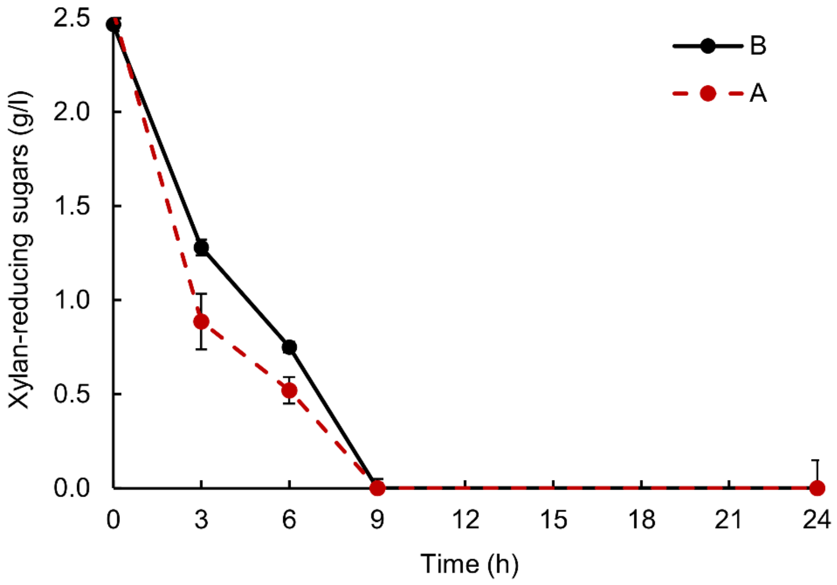

**Figure S2. Kinetics of xylan consumption in the bioreactors within 24 hours.** During the fluctuations at pH 6.0 (day 67) in reactor A, dense sampling within a 24 hours period showed that the fed water-soluble xylan was consumed within the first nine hours in both bioreactors. A and B stand for bioreactors A (pH 6.0) and B (pH 5.5). Error bars indicate the standard deviation (n=6).

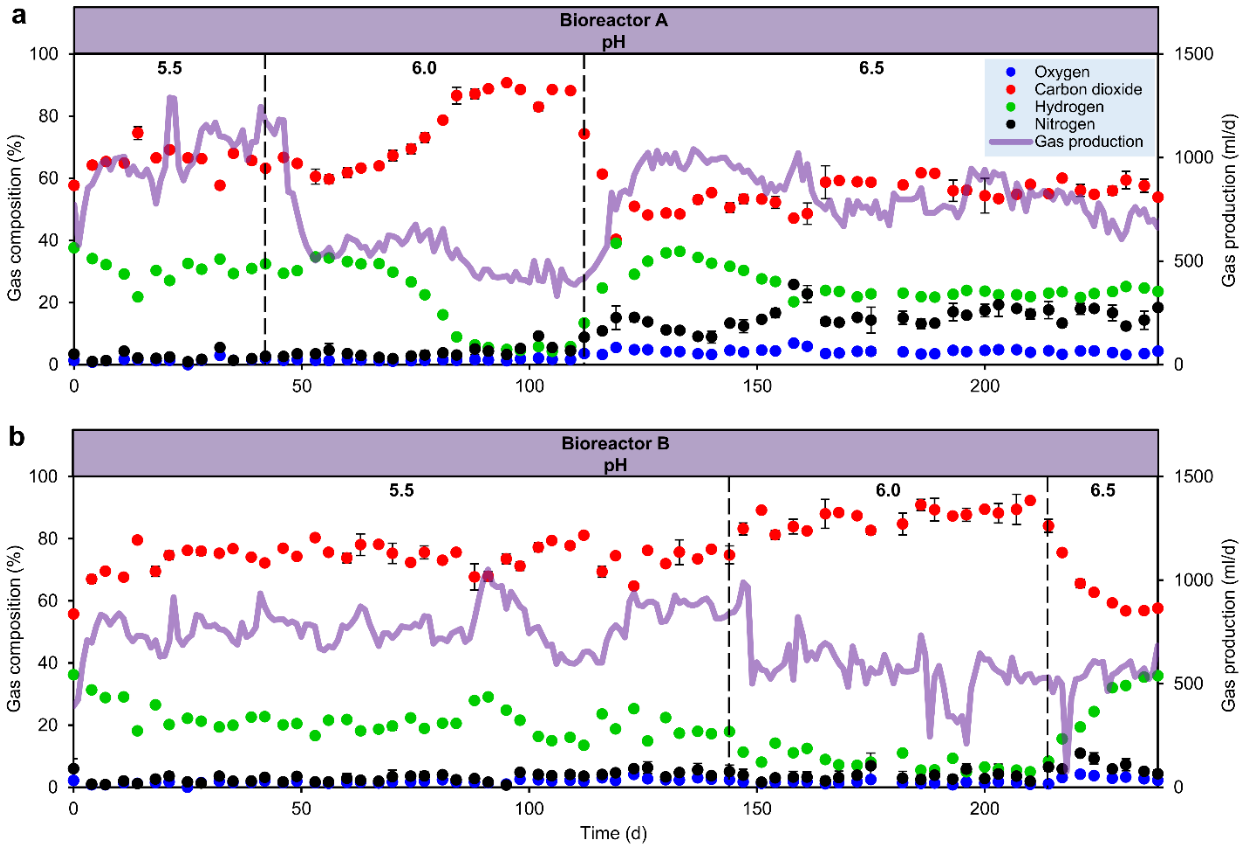

**Figure S3. Gas production of the bioreactors.** Daily gas production and gas composition in bioreactors A (**a**) and B (**b**) at three pH levels. Methane production was not detected. Error bars indicate the standard deviation (n=3).

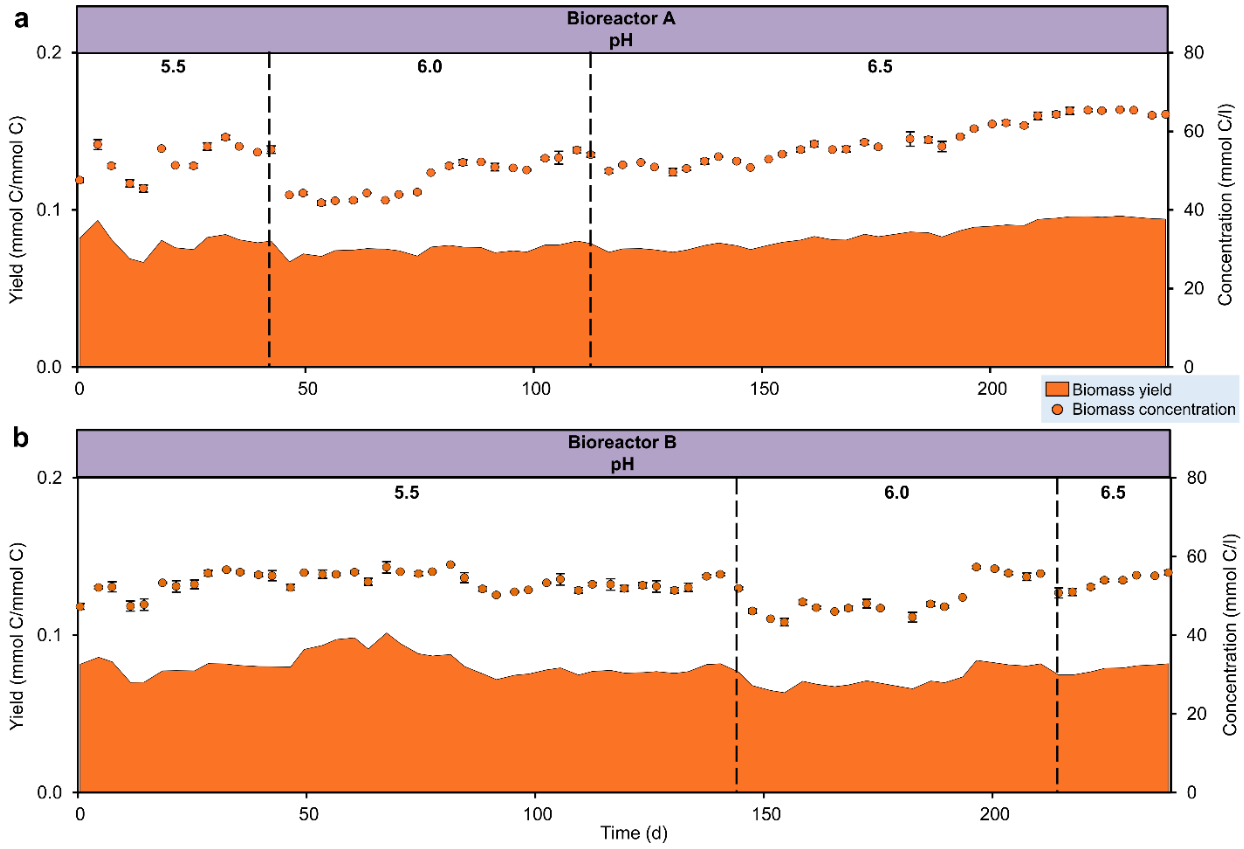

**Figure S4. Biomass production of the bioreactors.** Cell concentration and biomass yield in bioreactors A (**a**) and B (**b**) at three pH levels. The carbon amount of cell biomass was calculated by assuming an elemental biomass composition of $CH_{1.8}O_{0.5}N_{0.2}$ (molar mass = 24.6 g $mol^{-1}$). Error bars indicate the standard deviation (n=3).

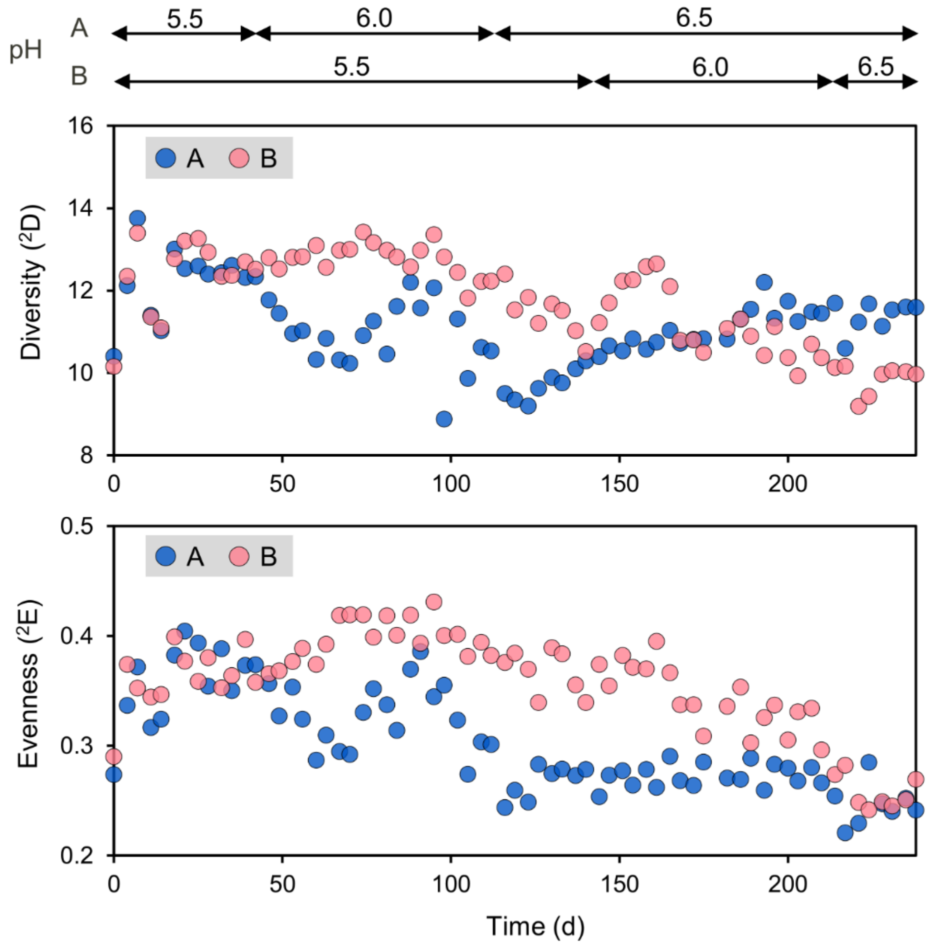

**Figure S5. Longitudinal changes in diversity and evenness of order two of bioreactor communities.** Based on the relative abundances of ASVs, α-diversity is represented by diversity of order two ($^2D$) and evenness of order two ($^2E$), which give more weight to the dominant types than to the rare types. A and B stand for bioreactors A and B.

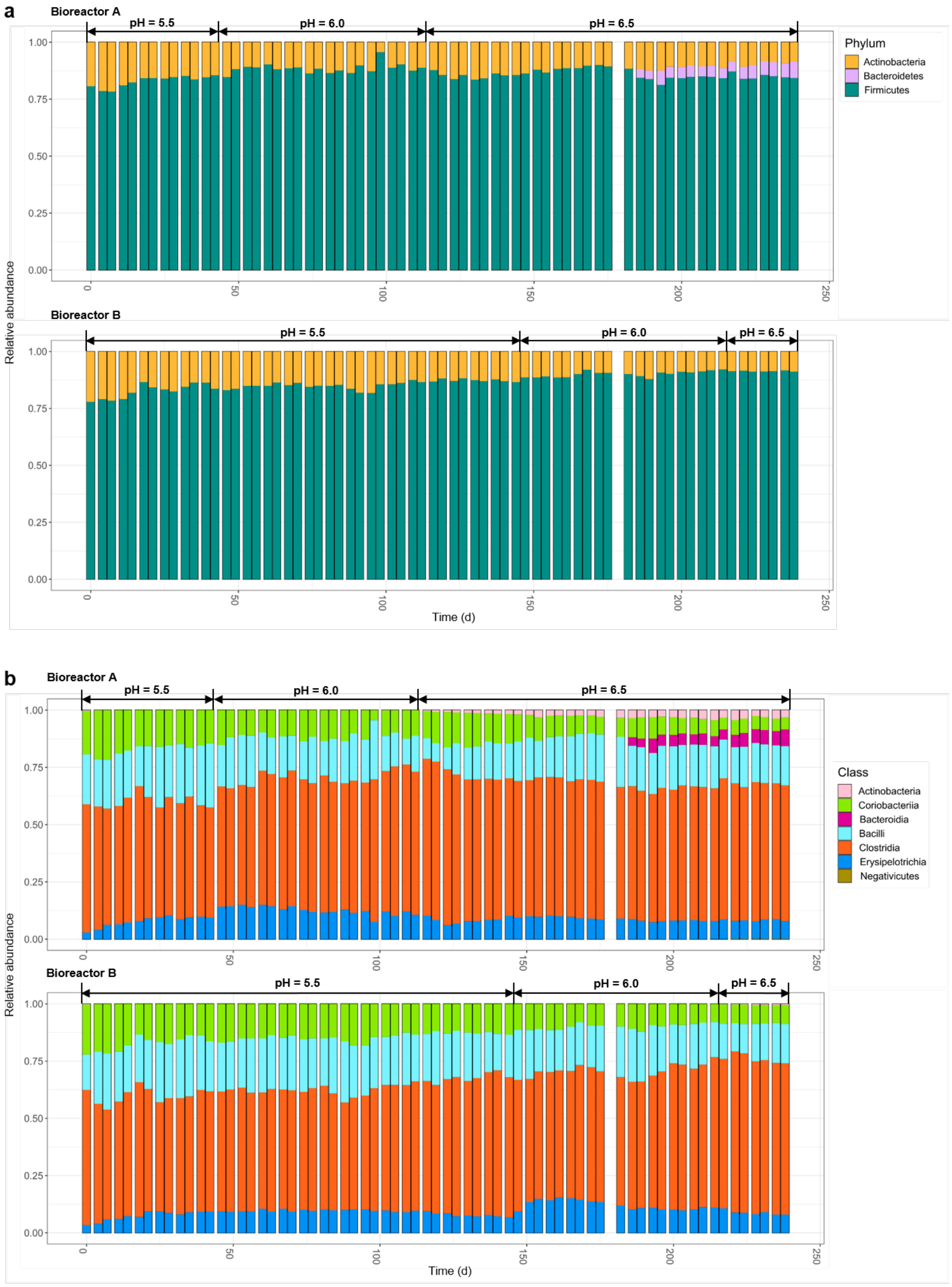
a
Bioreactor A
pH = 5.5
pH = 6.0
pH = 6.5
Bioreactor B
pH = 5.5
pH = 6.0
pH = 6.5
Relative abundance
Time (d)
Phylum
Actinobacteria
Bacteroidetes
Firmicutes
b
Bioreactor A
pH = 5.5
pH = 6.0
pH = 6.5
Bioreactor B
pH = 5.5
pH = 6.0
pH = 6.5
Relative abundance
Time (d)
Class
Actinobacteria
Coriobacteriia
Bacteroidia
Bacilli
Clostridia
Erysipelotrichia
Negativicutes

c

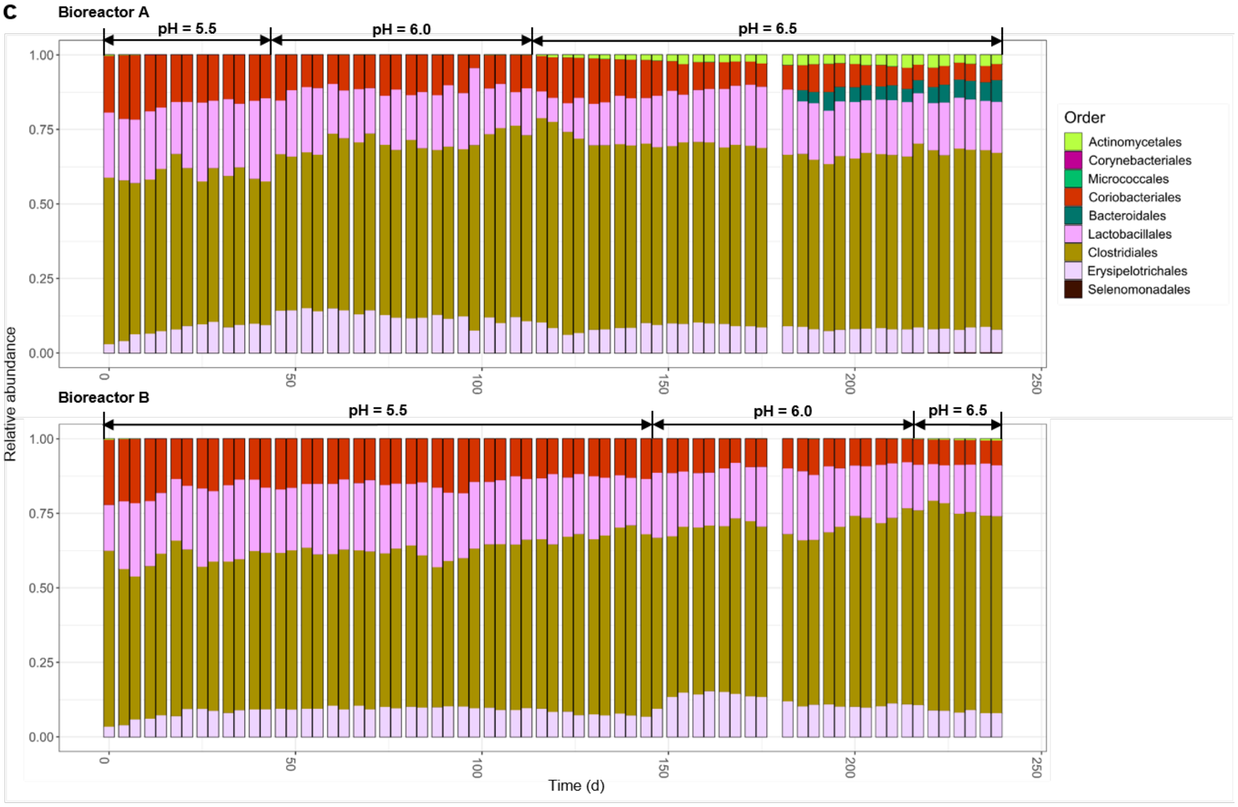

Bioreactor A
pH = 5.5
pH = 6.0
pH = 6.5
Bioreactor B
pH = 5.5
pH = 6.0
pH = 6.5
Relative abundance
Time (d)
Order
Actinomycetales
Corynebacteriales
Micrococcales
Coriobacteriales
Bacteroidales
Lactobacillales
Clostridiales
Erysipelotrichales
Selenomonadales

d

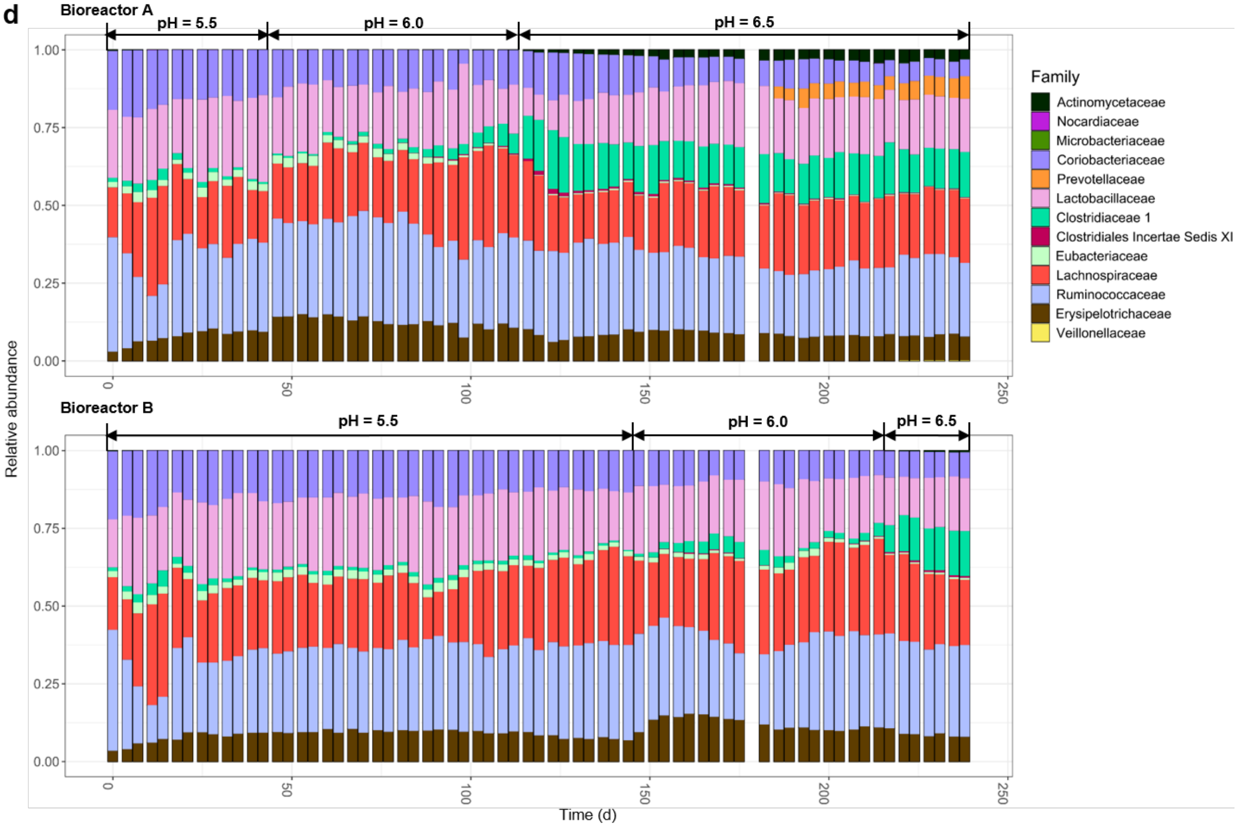

Bioreactor A
pH = 5.5
pH = 6.0
pH = 6.5
Bioreactor B
pH = 5.5
pH = 6.0
pH = 6.5
Relative abundance
Time (d)
Family
Actinomycetaceae
Nocardiaceae
Microbacteriaceae
Coriobacteriaceae
Prevotellaceae
Lactobacillaceae
Clostridiaceae 1
Clostridiales Incertae Sedis XI
Eubacteriaceae
Lachnospiraceae
Ruminococcaceae
Erysipelotrichaceae
Veillonellaceae

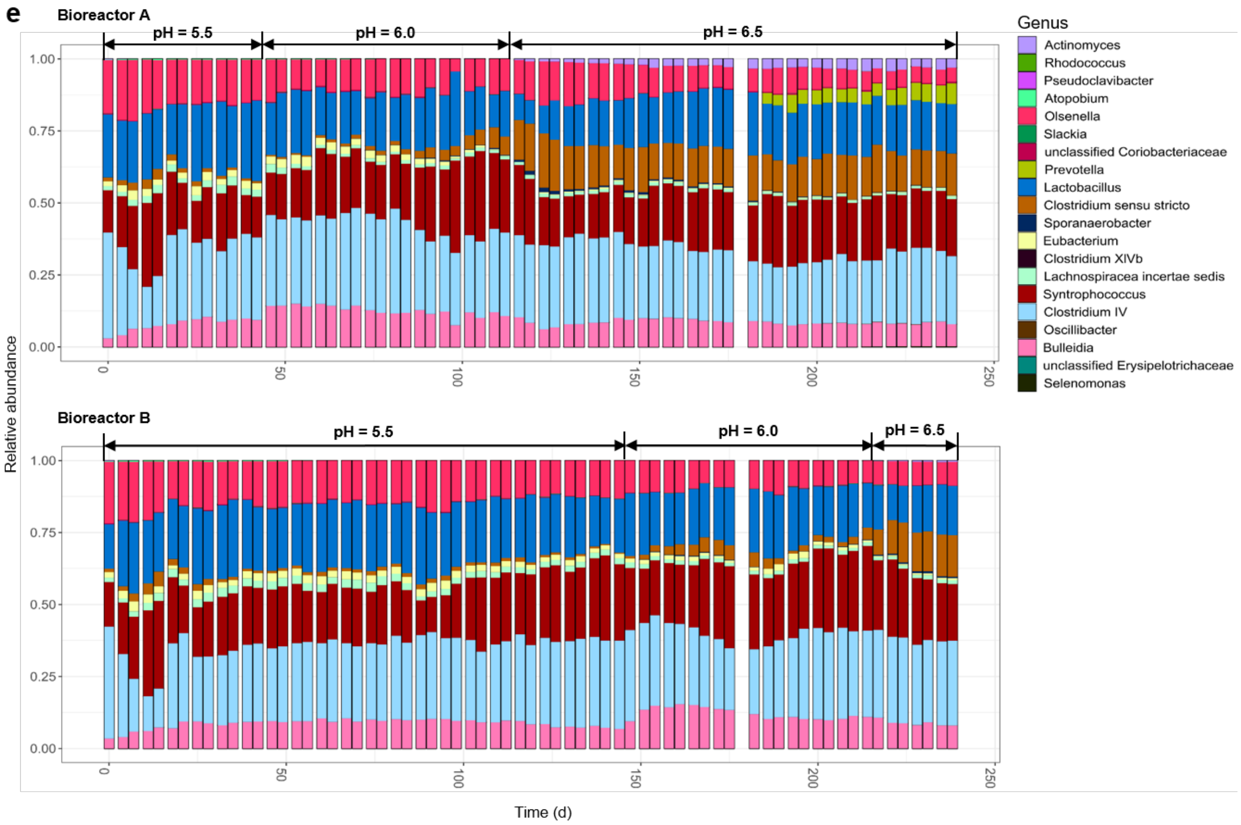

**Figure S6. Microbial community composition profiles of the bioreactors.** Based on amplicon sequencing of 16S rRNA genes, the taxonomic classification of amplicon sequence variants (ASVs) was categorized at the phylum (**a**), class (**b**), order (**c**), family (**d**) and genus (**e**) levels.

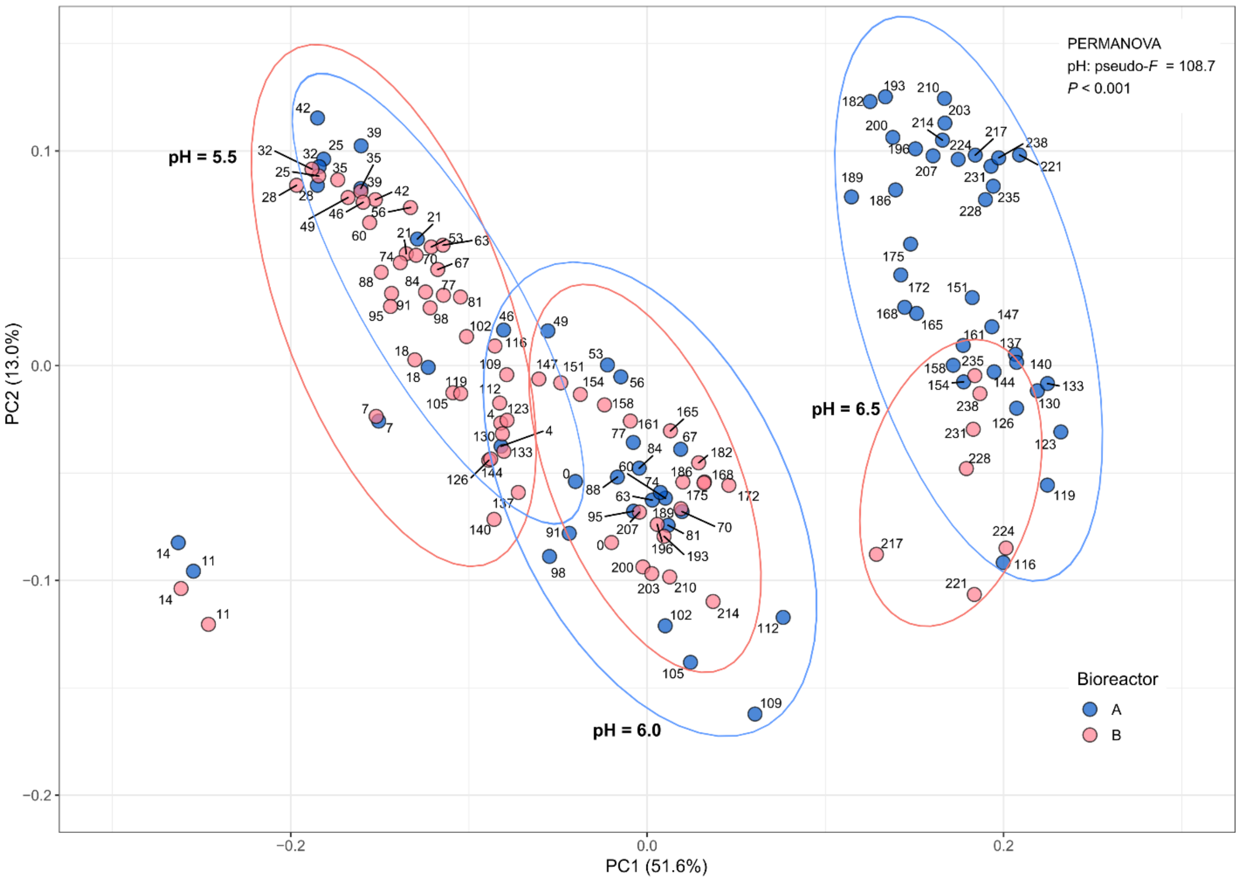

**Figure S7. Dissimilarities in bacterial community composition (β-diversity) of the two bioreactors A and B.** Principal coordinates analysis (PCoA) based on Bray-Curtis dissimilarities of microbial community composition in the bioreactors. Dots are named according to sampling days. Ellipses of 95% confidence intervals were added to each individual pH level of the bioreactors.

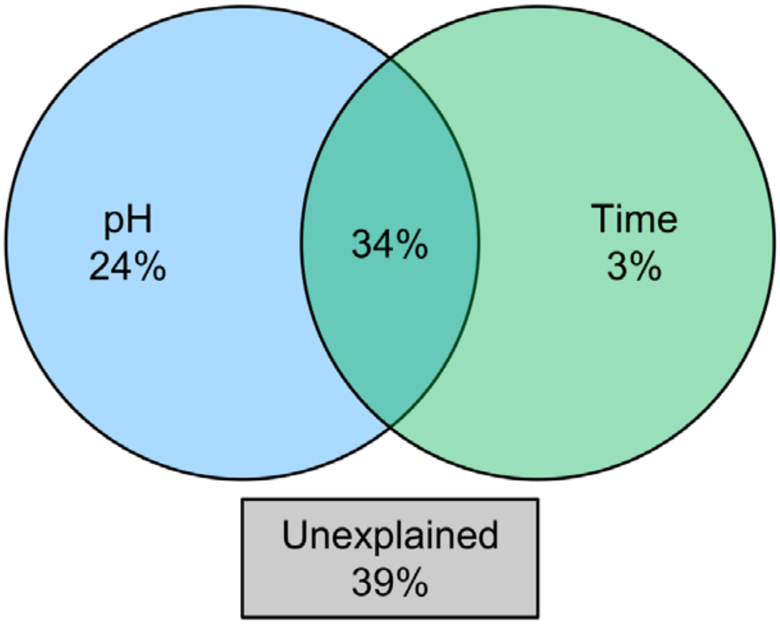

**Figure S8. Variation partitioning analysis (VPA) showing the relative importance of pH and time on microbial community variations.** VPA was used with redundancy analysis (RDA), and multiple partial RDAs were run to determine the partial, linear effect of each explanatory matrix in the response data. Numbers represent adjusted coefficients of determination (Adj. $R^2$ values).

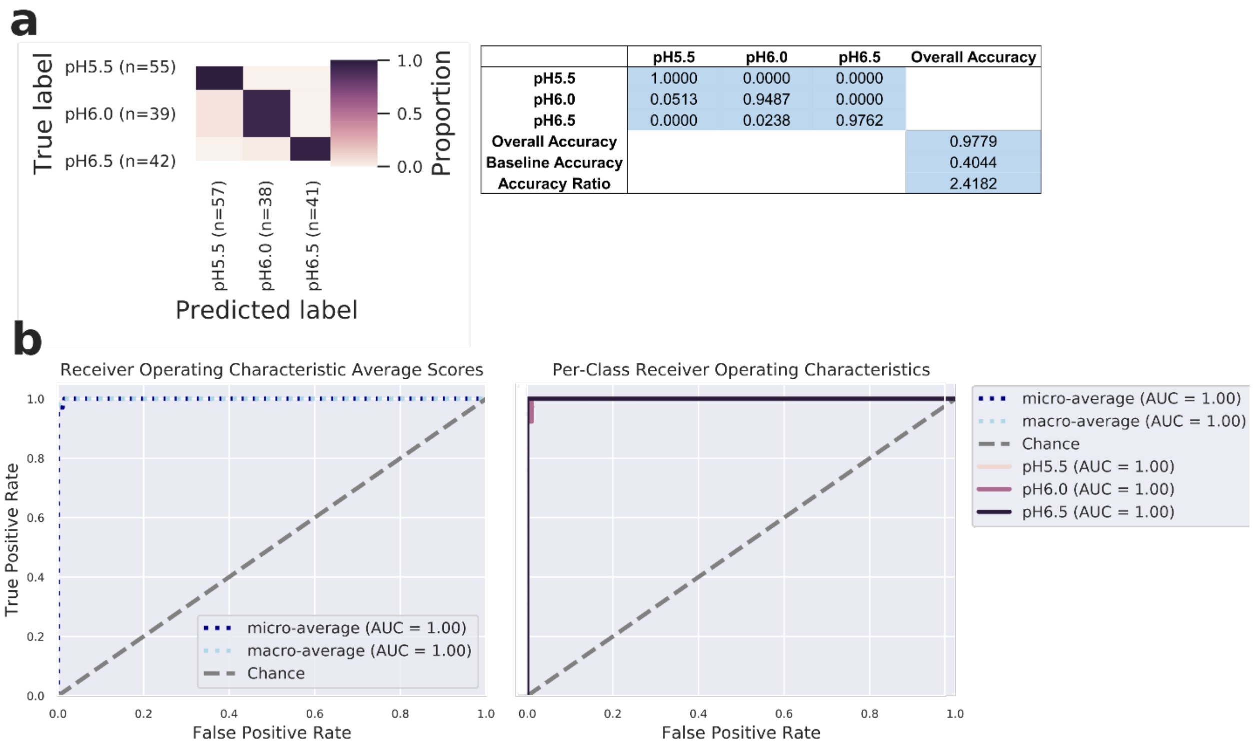

| | pH5.5 | pH6.0 | pH6.5 | Overall Accuracy |
|---|---|---|---|---|
| **pH5.5** | 1.0000 | 0.0000 | 0.0000 | |
| **pH6.0** | 0.0513 | 0.9487 | 0.0000 | |
| **pH6.5** | 0.0000 | 0.0238 | 0.9762 | |
| **Overall Accuracy** | | | | 0.9779 |
| **Baseline Accuracy** | | | | 0.4044 |
| **Accuracy Ratio** | | | | 2.4182 |

**Figure S9. Nested cross-validation of random forest classification in the prediction of pH levels for each sample. a,** Confusion matrix for the random forest classifier of all samples at three pH levels. For model optimization, two layers of *K*-fold ($K$ = 5) cross validation was incorporated to split the dataset into training and test set. Five different final models were trained, each sample received a predicted value and feature importance scores were averaged across each iteration. Overall accuracy was calculated by comparing the predicted values to the true values. **b,** The Receiver Operating Characteristic (ROC) and Area Under the Curve (AUC) curves represent the classification accuracy of the random forest. The ROC curve plots the relationship between the true positive rate and the false positive rate at various threshold settings. The AUC indicates the probability that the classifier ranks a randomly chosen sample of the given class higher than other classes. The random chance is represented as a diagonal line extending from the lower-left to the upper-right corner. In addition to show the ROC curves for each class, average ROCs and AUCs were calculated. "Micro-averaging" calculates metrics globally by averaging across each sample; hence class imbalance impacts this metric. "Macro-averaging" gives equal weight to the classification of each sample.

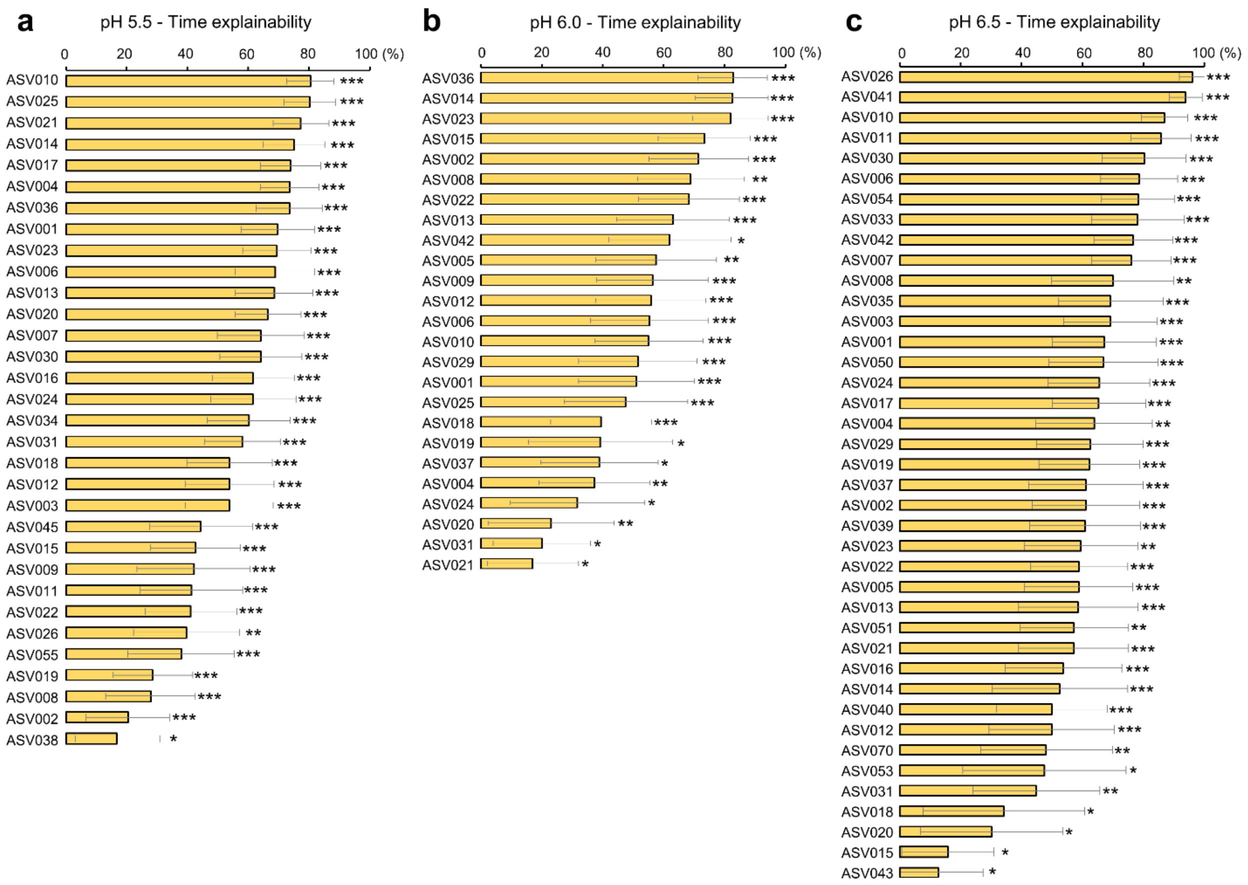

**Figure S10. Core time-dependent taxa of individual pH levels.** Using relative abundance data of ASVs of both bioreactors, a Microbial Temporal Variability Linear Mixed Model (MTV-LMM) was applied to identify time-dependent taxa of each individual pH level, whose abundance can be predicted based on the previous microbial community composition. As described, the time-explainability is denoted as the temporal variance explained by the microbial community in the previous time points. The time-explainability *P*-values: *P**** < 0.001 < ** < 0.01 < * < 0.05.

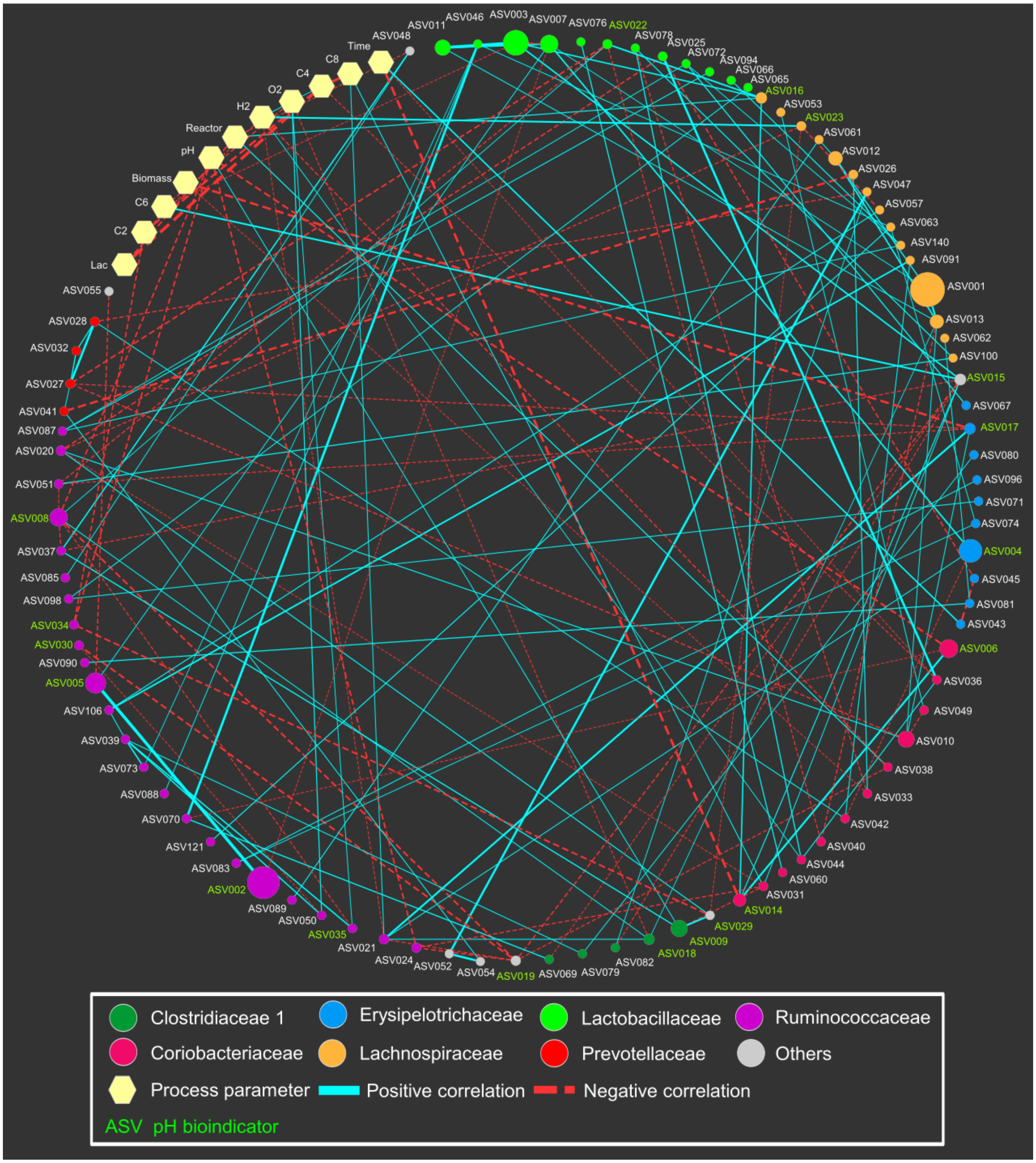

**Figure S11. Co-occurrence network for the entire period of reactor operation.** Edges indicate the significant ($P < 0.05$) correlations. Edge thickness reflects the strength of the correlation. Size of each ASV node is proportional to the mean relative abundance over the whole period. ASV nodes are colored and grouped by family. "Others" include the ASVs belonging to families *Eubacteriaceae* (ASV015),

*Actinomycetaceae* (ASV019), *Clostridiales incertae sedis* XI (ASV029), *Microbacteriaceae* (ASV048), *Veillonellaceae* (ASV052, ASV054) and *Nocardiaceae* (ASV055). pH bioindicators identified by random forest classification are shown with green letters. Lac, lactate concentration; C2, acetate yield; C4, *n*-butyrate yield; C6, *n*-caproate yield; C8, *n*-caprylate yield.

**Table S1. Linear mixed-effects model results for diversity of order one ($^1D$).** We considered time and pH as the fixed effects, and bioreactor as the random effect.

| Variable or parameter | Coefficient | Standard error | *Z*-score | *P* value |
|---|---|---|---|---|
| (Intercept) | 50.883 | 7.718 | 6.593 | < 0.001 |
| Time | -0.209 | 0.021 | -9.743 | < 0.001 |
| pH | -6.188 | 1.327 | -4.663 | < 0.001 |
| Time:pH | 0.035 | 0.004 | 9.777 | < 0.001 |
| Var. pH[a] | 0.396 | 1.071 | | |
| Var. Bioreactor [T.B][b] | 0.217 | | | |
| Cov. (pH, bioreactor)[c] | 0.953 | | | |

[a]Variance of pH

[b]Variance of bioreactor [treatment of bioreactor B]

[c]Covariance of pH and bioreactor (random intercept)

**Table S2. Linear mixed-effects model results for evenness of order one ($^1E$).** We considered time and pH as the fixed effects, and bioreactor as the random effect.

| Variable or parameter | Coefficient | Standard error | *Z*-score | *P* value |
|---|---|---|---|---|
| (Intercept) | 0.808 | 0.128 | 6.322 | < 0.001 |
| Time | 0.002 | 0.001 | 2.709 | 0.007 |
| pH | -0.061 | 0.022 | -2.720 | 0.007 |
| Time:pH | < -0.001 | < 0.001 | -2.741 | 0.006 |
| Var. pH[a] | < 0.001 | 0.002 | | |
| Var. Bioreactor [T.B][b] | < 0.001 | | | |
| Cov. (pH, bioreactor)[c] | < 0.001 | | | |

[a]Variance of pH

[b]Variance of bioreactor [treatment of bioreactor B]

[c]Covariance of pH and bioreactor (random intercept)

**Table S3. Linear mixed-effects model results for richness.** We considered time and pH as the fixed effects, and bioreactor as the random effect.

| Variable or parameter | Coefficient | Standard error | *Z*-score | *P* value |
|---|---|---|---|---|
| (Intercept) | 85.190 | 12.733 | 6.690 | < 0.001 |
| Time | 0.674 | 0.073 | 9.179 | < 0.001 |
| pH | 9.045 | 2.227 | 4.062 | < 0.001 |
| Time:pH | 0.116 | 0.012 | 9.344 | < 0.001 |
| Var. pH[a] | 0.466 | 0.620 | | |
| Var. Bioreactor [T.B][b] | 1.024 | | | |
| Cov. (pH, bioreactor)[c] | 13.738 | | | |

[a]Variance of pH

[b]Variance of bioreactor [treatment of bioreactor B]

[c]Covariance of pH and bioreactor (random intercept)

**Table S4. Linear mixed-effects model results for the relative abundance of *Clostridium* IV ASV008 at the different pH levels.** We considered time and pH as the fixed effects, and bioreactor as the random effect.

| Variable or parameter | Coefficient | Standard error | *Z*-score | *P* value |
|---|---|---|---|---|
| (Intercept) | 0.499 | 0.108 | 4.599 | < 0.001 |
| Time | 0.002 | 0.001 | 3.119 | 0.002 |
| pH | -0.077 | 0.019 | -4.123 | < 0.001 |
| Time:pH | < -0.001 | < 0.001 | -2.864 | 0.004 |
| Var. pH[a] | < 0.001 | 0.008 | | |
| Var. Bioreactor [T.B][b] | < -0.001 | | | |
| Cov. (pH, bioreactor)[c] | < 0.001 | | | |

[a]Variance of pH

[b]Variance of bioreactor [treatment of bioreactor B]

[c]Covariance of pH and bioreactor (random intercept)

**Table S5. Linear mixed-effects model results for the relative abundance of *Clostridium sensu stricto* ASV009 at the different pH levels.** We considered time and pH as the fixed effects, and bioreactor as the random effect.

| Variable or parameter | Coefficient | Standard error | *Z*-score | *P* value |
|---|---|---|---|---|
| (Intercept) | -0.081 | 0.220 | -4.003 | < 0.001 |
| Time | 0.001 | < 0.001 | 1.691 | 0.091 |
| pH | 0.156 | 0.037 | 4.252 | < 0.001 |
| Time:pH | < -0.001 | < 0.001 | -1.804 | 0.071 |
| Var. pH[a] | 0.002 | 0.114 | | |
| Var. Bioreactor [T.B][b] | < -0.001 | | | |
| Cov. (pH, bioreactor)[c] | < 0.001 | | | |

[a]Variance of pH

[b]Variance of bioreactor [treatment of bioreactor B]

[c]Covariance of pH and bioreactor (random intercept)

**Table S6. Linear mixed-effects model results for microbial community composition that is represented by the PC1 from the Aitchison distance-based principal component analysis.** We considered time and pH as the fixed effects, and bioreactor as the random effect.

| Variable or parameter | Coefficient | Standard error | *Z*-score | *P* value |
|---|---|---|---|---|
| (Intercept) | -1.033 | 0.254 | -4.059 | < 0.001 |
| Time | 0.001 | 0.001 | 1.683 | 0.092 |
| pH | 0.169 | 0.043 | 3.909 | < 0.001 |
| Time:pH | < -0.001 | < 0.001 | -1.387 | 0.165 |
| Var. pH[a] | 0.002 | 0.070 | | |
| Var. Bioreactor [T.B][b] | -0.001 | | | |
| Cov. (pH, bioreactor)[c] | < 0.001 | | | |

[a]Variance of pH

[b]Variance of bioreactor [treatment of bioreactor B]

[c]Covariance of pH and bioreactor (random intercept)

**Table S7. Linear mixed-effects model results for microbial community composition that is represented by the PC1 from the Bray-Curtis distance-based principal coordinate analysis.** We considered time and pH as the fixed effects, and bioreactor as the random effect.

| Variable or parameter | Coefficient | Standard error | *Z*-score | *P* value |
|---|---|---|---|---|
| (Intercept) | -2.216 | 0.235 | -9.428 | < 0.001 |
| Time | 0.004 | 0.001 | 3.161 | 0.002 |
| pH | 0.369 | 0.041 | 9.081 | < 0.001 |
| Time:pH | -0.001 | < 0.001 | -2.952 | 0.003 |
| Var. pH[a] | < 0.001 | 0.014 | | |
| Var. Bioreactor [T.B][b] | < -0.001 | | | |
| Cov. (pH, bioreactor)[c] | 0.001 | | | |

[a]Variance of pH

[b]Variance of bioreactor [treatment of bioreactor B]

[c]Covariance of pH and bioreactor (random intercept)

**Table S8. Summary statistics of networks.**

| Dataset | No. of Nodes | No. of Edges | avgN[a] | avgCC[b] | Density | Heterogeneity | Centralisation |
|---|---|---|---|---|---|---|---|
| Entire | 100 | 151 | 3.256 | 0.062 | 0.038 | 0.469 | 0.057 |
| pH 5.5 | 70 | 77 | 2.612 | 0.039 | 0.054 | 0.512 | 0.074 |
| pH 6.0 | 60 | 63 | 2.455 | 0.029 | 0.057 | 0.447 | 0.062 |
| pH 6.5 | 86 | 99 | 2.528 | 0.078 | 0.036 | 0.559 | 0.065 |

[a]avgN, average number of neighbours

[b]avgCC, average clustering coefficient

**Table S9. Metagenome-assembled genomes (MAGs) with the same taxonomy as ASVs.**

| ASV | Taxonomic classification[a] | | | | | | Representative MAG | Accession number[b] |
|---|---|---|---|---|---|---|---|---|
| | Phylum | Class | Order | Family | Genus | Species | | |
| *Clostridium* IV sp. ASV008 | *Firmicutes_A* | *Clostridia* | *Oscillospirales* | *Acutalibacteraceae* | UBA1033 | UBA1033 sp002399935 | UMB00014 | GCA_903789645 |
| | *Firmicutes_A* | *Clostridia* | *Oscillospirales* | *Acutalibacteraceae* | UBA1033 | UBA1033 sp002407675 | UMB00060 | GCA_903789675 |
| | *Firmicutes_A* | *Clostridia* | *Oscillospirales* | *Acutalibacteraceae* | UBA1033 | UBA1033 sp002409675 | UMB00097 | GCA_903789585 |
| | *Firmicutes_A* | *Clostridia* | *Oscillospirales* | *Acutalibacteraceae* | UBA4871 | UBA4871 sp002399445 | UMB00016 | GCA_903789565 |
| *Clostridium sensu stricto* sp. ASV009 | *Firmicutes_A* | *Clostridia* | *Clostridiales* | *Clostridiaceae* | *Clostridium_B* | *Clostridium_B* sp003497125 | UMB00080 | GCA_903789665 |
| *Olsenella* sp. ASV049 | *Actinobacteriota* | *Coriobacteriia* | *Coriobacteriales* | *Atopobiaceae* | *Olsenella_B* | *Olsenella_B* sp000752675 | UMB00010 | GCA_903789475 |
| | *Actinobacteriota* | *Coriobacteriia* | *Coriobacteriales* | *Atopobiaceae* | *Olsenella_C* | unclassified | UMB00003 | GCA_903789455 |
| | *Actinobacteriota* | *Coriobacteriia* | *Coriobacteriales* | *Atopobiaceae* | *Olsenella* | unclassified | UMB00074 | GCA_903789485 |
| *Lactobacillus* spp. ASV046, ASV065, ASV076 | *Firmicutes* | *Bacilli* | *Lactobacillales* | *Lactobacillaceae* | *Lactobacillus_H* | *Lactobacillus_H mucosae* | UMB00017 | GCA_903789575 |
| | *Firmicutes* | *Bacilli* | *Lactobacillales* | *Lactobacillaceae* | *Lactobacillus* | unclassified | UMB00041 | GCA_903789695 |
| | *Firmicutes* | *Bacilli* | *Lactobacillales* | *Lactobacillaceae* | *Lactobacillus* | *Lactobacillus amylovorus* | UMB00015 | GCA_903789705 |

[a] Taxonomy refers to the Genome Taxonomy Database (GTDB) phylogenomic classification

[b] Accession numbers refer to the European Nucleotide Archive (ENA)
